# Identification of AB8939, a novel synthetic microtubule destabilizer and ALDH inhibitor that overcomes multidrug resistance in tumor cells as a drug candidate for the treatment of refractory acute myeloid leukemia

**DOI:** 10.64898/2025.12.10.692519

**Authors:** Martine Humbert, Sébastien Letard, Armelle Goubard, Camille Montersino, Stéphane Audebert, Emilie Baudelet, Bérengère Hajem, Manuel Neves, Samila Siavioshian, Paloma Fernandez-Varela, Benoît Gigant, Etienne Rebuffet, Pascal Verdier-Pinard, Rémy Castellano, Yves Collette, Norbert Vey, Abdellah Benjahad, Didier Pez, Jason Martin, Alain Moussy, Colin Mansfield, Laurent Gros, Christian Auclair, Patrice Dubreuil

**Affiliations:** AB Science, Paris, France; INSERM, CNRS, Institut Paoli-Calmettes, CRCM, Aix-Marseille Univ., 27 Boulevard Lei Roure, 13009 Marseille, France; Paris-Saclay University, CEA, CNRS, Institute for Integrative Biology of the Cell (I2BC), Gif-sur-Yvette, France; INSERM, CNRS, Institut Paoli-Calmettes, CRCM, Marseille Protéomique, Aix-Marseille University, 27 Boulevard Lei Roure, 13009 Marseille, France; Aix-Marseille University, Inserm, CNRS, Institut Paoli-Calmettes, CRCM, TrGET plateform, 13009 Marseille, France; Aix Marseille Université, CNRS, INP UMR7051, NeuroCyto, 13005, Marseille, France

## Abstract

We identified AB8939, a novel small synthetic molecule that exhibits strong and broad antiproliferative activity against a panel of various cancer cell types with IC_50_ values in the nanomolar range. In vitro investigations showed that AB8939 is a novel microtubule-targeting agent that interacts with the colchicine-binding site of tubulin. AB8939 disrupts the microtubule network, leading to mitotic arrest in G2/M phase and subsequent apoptosis. Importantly, AB8939 overcomes drug resistance mechanisms, including overexpression of efflux transporters such as P-glycoprotein (P-gp) and aberrant expression of β3-tubulin. AB8939 displays high cytotoxicity against blasts from AML patients, including blasts resistant to cytarabine (Ara-C). In vivo, AB8939 shows strong antitumor activity in MOLM-14, an Ara-C-resistant AML model, as evidenced by tumor growth inhibition and substantial increase in mouse survival. Further experiments performed on an AML PDX TG-AML-36 model demonstrated that AB8939 efficiently kills leukemic stem cells (CD34+/CD38−). Reverse proteomic experiments revealed that AB8939 inhibits ALDH1 and ALDH2, enzymes often overexpressed in tumors and tumor stem cells, thereby favoring tumor progression and relapse. AB8939 is a novel dual-targeting drug that acts on both tubulin and ALDH enzymes, with potential activity against various cancer types, especially refractory AML with complex karyotypes such as those displaying *MECOM* rearrangement and AML with mutations associated with poor prognosis, such as *ASXL1* and *TP53*.

## INTRODUCTION

Acute myeloid leukemia (AML) is the most common form of acute leukemia in adults and is commonly accompanied by a poor prognosis, with less than 25% of patients surviving five years after diagnosis (1). The genetic and phenotypic heterogeneity of AML complicates treatment strategy, as the disease encompasses a variety of genetic mutations and epigenetic modifications that influence tumor behavior and response to therapy (2). Mutations in genes such as *FLT3*, *NPM1*, and *IDH1/2* can significantly alter treatment outcomes and resistance profiles. Patient age and the presence of comorbid conditions can impact the ability to tolerate intensive therapies. Additionally, the patient’s cytogenetic profile is critical; complex karyotypes or deletions of chromosomes 5 or 7 are associated with lower survival rates and reduced response to standard therapies. Finally, minimal residual disease (MRD)—the presence of residual leukemic stem cells after therapy—can complicate further treatment options and effectiveness. Treatment-related complications, drug resistance, and the potential for disease relapse further complicate the effective management of AML, highlighting the need for new therapeutic approaches.

As part of a program focused on discovering novel small cytotoxic molecules capable of overcoming drug resistance, we conducted a phenotypic cell-based antiproliferative screen of the AB Science small molecule library using a panel of approximately 80 human cancer cell lines representing 17 cancer types. From this screen, we identified AB8939, a small chemical molecule with strong and broad antiproliferative activity showing high similarity to microtubule-targeting agents, including vincristine and colchicine. In vitro studies confirmed that AB8939 inhibited polymerization of purified porcine tubulin, similar to that reported for known microtubule polymerization inhibitors (colchicine, vincristine, VERU-111). Further investigations using X-ray crystallography showed that AB8939 binds to the colchicine site of β-tubulin. Studies using several cancer cell lines demonstrated that AB8939 caused rapid and profound disruption of the cellular microtubule network and led to cell cycle arrest at the G2/M phase followed by apoptosis.

Importantly, AB8939 was shown to overcome major drug resistance mechanisms. Ex vivo data demonstrate that AB8939 displays marked cytotoxic effects in cell lines expressing P-glycoprotein (P-gp) and resistant to various other antitumor agents, including doxorubicin and cytarabine (Ara-C). Furthermore, AB8939 is effective ex vivo on blasts isolated from both newly diagnosed (ND) and relapsed or refractory (R/R) AML patients, the latter being associated with extremely poor outcomes and limited therapeutic options (Thol et al., 2020).

In vivo data from an Ara-C-resistant mouse MOLM-14 model of AML showed that AB8939 administered alone is non-toxic to mice and has a potent antitumor effect, decreasing tumor growth and increasing survival. A second set of experiments using the AML PDX TG-AML-36 model showed that AB8939 strongly decreases tumor recurrence, suggesting that the drug targets leukemic stem cells as well.

To identify other possible pharmacological targets that could account for the effect of AB8939 on tumor recurrence, we carried out a reverse proteomic experiment that demonstrated AB8939’s ability to bind to aldehyde dehydrogenases (ALDH1s and ALDH2). AB8939 acts as a potent inhibitor of ALDH1A1 and ALDH2 using recombinant enzymes. High expression levels of these enzymes are associated with enhanced cell growth, increased motility, and improved invasion capabilities of cancer cells across various cancer types, including breast, lung, colorectal, and prostate cancers (3). High levels of ALDH2 are often associated with enhanced cancer cell survival, proliferation, and resistance to conventional therapies. ALDH2 contributes to several critical processes in tumors, including maintaining a stem-cell-like phenotype, promoting epithelial-mesenchymal transition (EMT), and regulating signaling pathways linked to tumorigenesis and metastasis (4).

This dual mechanism of action allows AB8939 to target both proliferative tumor cells and quiescent leukemic stem cells. This remarkable property makes AB8939 an appropriate drug candidate for the treatment of refractory AML.

## RESULTS

### AB8939 Shows Strong and Broad Antiproliferative Activity In Vitro

AB8939 is a small synthetic molecule that was initially synthesized as part of a protein kinase drug discovery program. Although this compound exhibited no kinase activity whatsoever in numerous in vitro kinase assays and KinomeScan analyses (results not shown), a phenotypic screening campaign performed against a panel of 33 cancer cell lines representative of various tumors (see Supplementary data, Table S1) aimed at identifying potent anticancer drugs in AB Science’s entire proprietary chemical library identified AB8939 as the most promising drug candidate for targeting hematological cancers. The chemical structure of AB8939 is presented in Figure S1 (Supplementary Data).

The in vitro cytotoxicity of AB8939 was assayed by determining its ability to inhibit proliferation of tumor cells against an extended panel of 83 cancer cell lines (Table 1). These in vitro data demonstrate excellent antiproliferative activity of AB8939. AB8939 inhibited cancer cell growth across a variety of cancer types and was particularly active against hematopoietic cancers. As shown in Table 1, more than 50% of tumor cell lines tested showed reduced proliferation/survival with an IC_50_ ≤500 nM following 72 hours of treatment, and most hematopoietic cell lines showed an IC_50_ ≤100 nM. For cell lines with IC_50_ ≥10 µM after 72 hours, they were shown to be highly sensitive to AB8939 when treated for extended incubation times (6 days) with IC_50_ <100 nM (Supplementary data, Table S1,); 16 out of 19 epithelial cell lines tested had IC_50_ ≤10 nM. Therefore, among the different human tumor cell lines tested, all were efficiently inhibited, including tumor cells with oncogenic driver mutations.

**Table 1.** Cytotoxicity profile (IC_50_ nM) of AB8939 against a panel of 83 human cancer cell lines and primary cells at 72h drug exposure. (CNS: Central Nervous System).

| Haematopoietic Tumor cell lines |  |  |  |  |  |  |  |
| --- | --- | --- | --- | --- | --- | --- | --- |
| **AML** | **HL-60** | **NOMO-1** | **THP-1** | **RS4-11** | **U-937** | **MV-4-11** | **MOLM-14** |
|  | 30 | 66 | 46 | 0.1 | 19 | 10 | 20 |
| **B cell Lymphoma** | **HLB-2** | **JVM-3** | **RAJI** | **REC-1** | **RL** | **SUDHL-4** | **RAMOS** |
|  | 10 | 1 | 10 | 4 | 10 | 10 | 10 |
| **T cell Lymphoma** | **CCRFCEM** | **HUT78** | **JURKAT** | **MOLT-13** | **RPMI8401** | **PEER** | **KARPAS299** |
|  | 1 | 4 | 13 | 9 | 35 | 9 | 1 |
| **Multiple Myeloma** | **OPM-2** | **RPMI8226** |  |  |  |  |  |
|  | 10 | 10 |  |  |  |  |  |
| Solid Tumor cell lines |  |  |  |  |  |  |  |
| **H&N** | **A64CLS** | **CLS354-4** | **Detroit562** | **Hep2** | **RPMI2650** | **SW579** |  |
|  | 10 | 10 | 10 | 10 | >10000 | 10 |  |
| **Melanoma** | **A375** | **LU1205** | **M6** | **M67** | **MELSOL** | **MELWO** | **SKMEL28** |
|  | 13 | 16 | >10000 | >10000 | 19 | 31 | >10000 |
| **Pancreas** | **BxPC-3** | **CAPAN-2** | **MIAPACA-2** | **PANC-1** |  |  |  |
|  | 10 | >10000 | >10000 | >10000 |  |  |  |
| **Liver** | **HEPG2** | **PLC PRF5** |  |  |  |  |  |
|  | >10000 | >10000 |  |  |  |  |  |
| **Stomach** | **AGS** | **HGC27** |  |  |  |  |  |
|  | 3995 | 624 |  |  |  |  |  |
| **Colon** | **DLD-1** | **HCT116** | **CACO-2** | **HRT-18** | **HT29** | **LS174T** | **SW480** |
|  | 10 | 10 | 176 | >10000 | 10 | >10000 | >10000 |
| **Lung** | **A549** | **CALU-6** | **GCT** | **H1299** | **H1650** | **H820** | **HCC827** |
|  | >10000 | 10 | 10 | >10000 | >10000 | 8960 | 2155 |
| **Breast** | **BT20** | **BT474** | **MCF-7** | **HCC1937** | **MDA-MB-134** | **MDA-MB-231** | **BRCA MZ01** |
|  | >10000 | >10000 | >10000 | 530 | >10000 | >10000 | >10000 |
| **Ovaries** | **MESSA** | **OV90** | **OVCAR3** | **TOV112D** | **TOV21G** | **O114** |  |
|  | >10000 | 3431 | 91 | 298 | 51 | 4821 |  |
| **Prostate** | **DU145** | **LnCAP** | **PC-3** |  |  |  |  |
|  | >10000 | >10000 | 10 |  |  |  |  |
| **Kidney** | **786-O** | **A498** | **ACHN** | **CAKI-1** | **CAKI-2** |  |  |
|  | 20 | >10000 | >10000 | >10000 | >10000 |  |  |
| CNS Tumors cell lines |  |  |  |  |  |  |  |
| **Neuroblastoma** | **SHSY5Y** |  | **Neuroglioma** | **H4** | **Glioblastoma** | **U118** | **U87MG** |
|  | 10 |  |  | 6 |  | 298 | >10000 |
| Primary cells |  |  |  |  |  |  |  |
| **Cardio-myocytes** |  | **Human Dermal Fibroblasts** |  |  |  |  |  |
| >10000 |  | >10000 |  |  |  |  |  |

### AB8939 Displays High Antiproliferative Activity on Acute Megakaryoblastic Leukemia (AMKL) Cells

Acute megakaryoblastic leukemia (AMKL) is a rare and aggressive subtype of acute myeloid leukemia characterized by the uncontrolled proliferation of malignant megakaryoblasts in the bone marrow (11). It predominantly affects children, particularly those with Down syndrome (DS). Current chemotherapy regimens for AMKL mirror those used in AML and typically include cytarabine and daunorubicin. However, both agents are substrates of P-glycoprotein (P-gp), which can contribute to the development of drug resistance. Clinically, DS children with AMKL exhibit high response rates and favorable 1Illyear survival, whereas nonIllDS AMKL is associated with greater malignancy, early relapse, and poor prognosis—underscoring the urgent need for improved therapeutic options in this population.

To address this, we compared the antiproliferative activity of AB8939 with doxorubicin in two megakaryoblastic cell lines, CHRF and CMK. CHRF cells express platelet peroxidase, platelet factor 4, platelet glycoprotein IIb–IIIa (CD41), factor VIII antigen, and the myeloid markers CD13 (MY7) and CD33 (MY9), as well as the transcription factor GATAIll1. CMK cells were established from a DS patient with AMKL and are positive for antiplatelet antibodies. In these cells, platelet peroxidase reactivity localizes to the nuclear envelope and endoplasmic reticulum. Cytogenetic analysis revealed nearIlltetraploidy and a der(17)t(11;17) translocation, also present in the patient’s original leukemic blasts, confirming their malignant origin.

AB8939 demonstrated markedly greater antiproliferative activity than doxorubicin in both models (Table 2). In CHRF cells, the IC₅₀ of AB8939 was 9.7 nM compared with 681 nM for doxorubicin, reflecting a >70Illfold increase in potency. In CMK cells, AB8939 achieved an IC₅₀ of 5.68 nM, whereas doxorubicin required 844 nM, corresponding to nearly a 150Illfold difference. These results indicate that AB8939 is consistently and substantially more potent than doxorubicin across distinct AMKL cellular contexts, including both nonIllDS (CHRF) and DSIllderived (CMK) backgrounds, highlighting its potential to overcome the limitations of conventional anthracyclines in this disease.

**Table 2.** Comparative antiproliferative effect of AB8939 and doxorubicin on two AMKL cell lines. Cells were incubated for 72 hours with either AB8939 or doxorubicin. (IC_50_ values are expressed in nM).

| Cell line | AB8939 (IC<sub>50</sub> nM) | Doxorubicin (IC<sub>50</sub> nM) |
| --- | --- | --- |
| CHRF | 9.70 | 681 |
| CMK | 5.68 | 844 |

These findings underscore the promise of AB8939 as a therapeutic alternative to conventional anthracyclines in AMKL. The striking potency advantage—over 70-fold in CHRF cells and nearly 150-fold in CMK cells—suggests that AB8939 retains activity in both non-DS and DS-derived AMKL contexts, where resistance to standard agents such as doxorubicin remains a major clinical challenge. The ability of AB8939 to achieve nanomolar efficacy in vitro further supports its potential to circumvent P-gp–mediated efflux and other mechanisms of anthracycline resistance, particularly relevant in non-DS AMKL where outcomes remain dismal. Collectively, these results provide a strong rationale for advancing AB8939 into further preclinical and clinical evaluation as a candidate therapy for high-risk AMKL populations.

### AB8939 Displays an Activity Profile Reminiscent of Tubulin-Destabilizing Agents

An in silico comparison of AB8939’s activity profile against the extended 83-cell line panel with several known cytotoxic agents having different mechanisms of action suggested that AB8939 may be acting as a microtubule-targeting agent (MTA). The similarity relative to AB8939 was expressed using Pearson’s correlation coefficient (PCC); a PCC value >0.5 is generally considered significant. As presented in Table 3, IC_50_ values of AB8939 were highly similar to most MTAs on hematopoietic cell lines following 72-hour assays, particularly with colchicine-binding site inhibitors and vincristine.

**Table 3.** Comparison of growth inhibitor patterns of AB8939 and 8 MTAs on 33 tumor cell lines in a 72 hours proliferation/survival assay, using the Pearson’s correlation coefficient (PPC).

| IC<sub>50</sub> (nM) | Colchicine Binding Site Inhibitors (CBSI) |  |  |  |  |  | Vinca-Alkaloid Binding Site |  | Taxane |
| --- | --- | --- | --- | --- | --- | --- | --- | --- | --- |
|  | AB8939 | Sabizabulin (Veru-111) | Unesbulin (PTC-596) | Colchicine | CA-4 | Lexibulin (CYT997) | Vincristine | Eribulin | Taxol |
| Average all cell lines | 3890 | 411 | 3770 | 6770 | 3691 | 3976 | 5644 | 5487 | 11576 |
| Average Hemato | 50 | 50 | 50 | 50 | 50 | 50 | 100 | 132 | 1382 |
| PCC All | 1.00 | 0.25 | 0.25 | 0.45 | 0.45 | 0.56 | 0.34 | 0.135 | 0.26 |
| PCC hemato | 1.00 | 0.87 | 0.84 | 0.78 | 0.91 | 0.70 | 0.67 | 0.13 | 0.035 |
| PCC non hemato | 1.00 | 0.23 | 0.16 | 0.33 | 0.38 | 0.5 | 0.21 | 0.00 | 0.22 |

*PCC=Pearson's Correlation coefficient, Hemato= Hematopoietic cell lines*

### AB8939 Induces Disruption of the Microtubule Network in Intact Cells

To assess the potential effect of AB8939 on cytoskeleton components, we first performed immunofluorescent staining of microtubules and actin filaments. The NIH 3T3 cell line was chosen for its large size and flat morphology. NIH 3T3 cells were treated for 1 hour with 100 nM AB8939, then processed for immunofluorescence staining for microtubules (anti-α-tubulin) and actin (phalloidin-FITC).

Figure 1A shows that AB8939 induces rapid and drastic disruption of microtubules while not affecting the actin network. Microtubule disassembly is observed within 1 hour of exposure to 100 nM AB8939. Similar microtubule disorganization was observed using different human tumor cell lines (data not shown).

**Figure 1.**
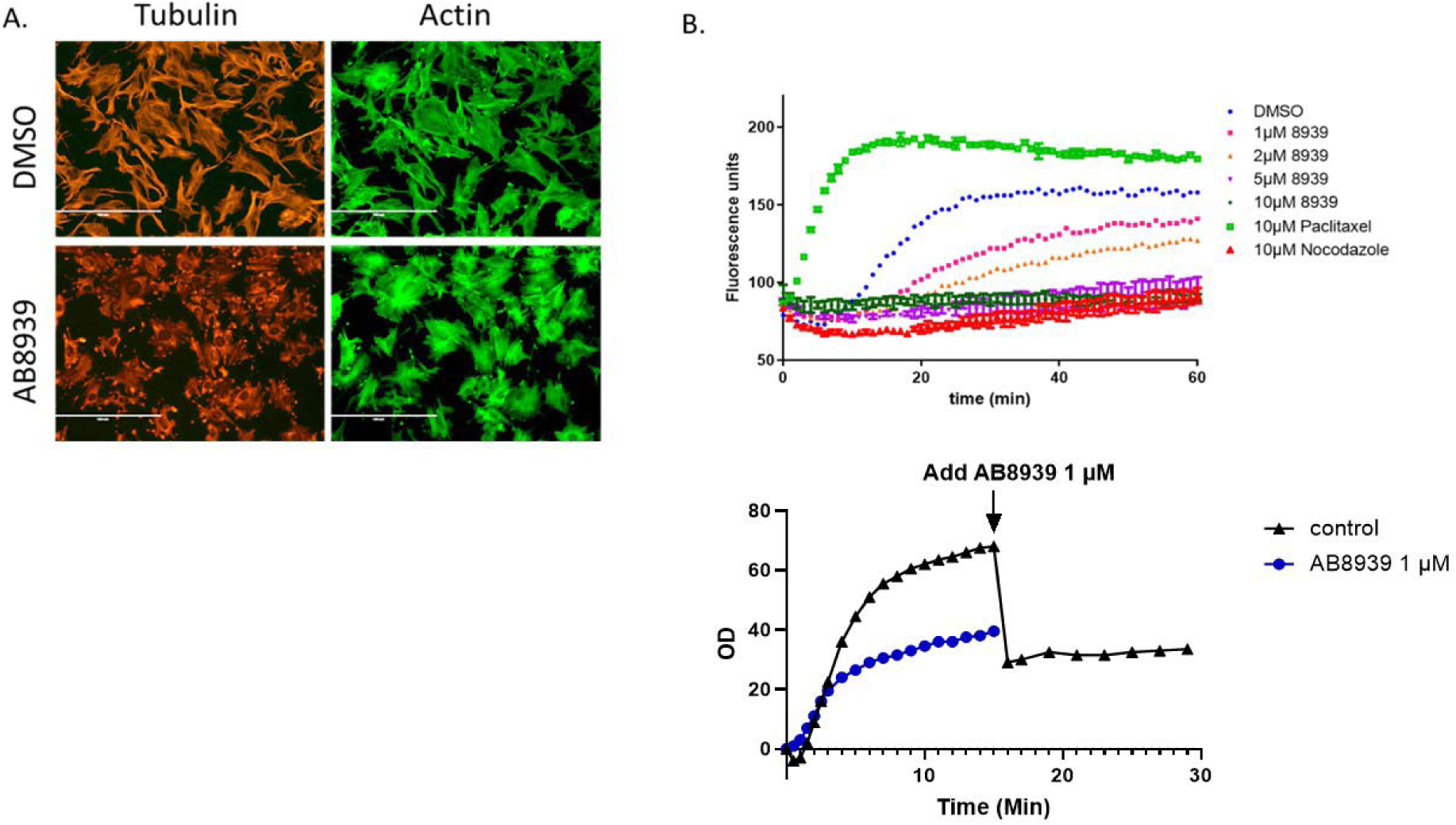
AB8939 disrupts the MT network and inhibits tubulin polymerization. Panel A: Immunofluorescent staining of microtubules (anti-α-tubulin, red) and actin (phalloidin-FITC, green) in NIH 3T3 cells treated with vehicle (DMSO) or 100 nM AB8939 for 1 hour. AB8939 rapidly disrupts the microtubule network while leaving actin intact. Scale bar = 20 µm. Panel B: Cell-free tubulin polymerization assay. Porcine brain tubulin (20 µM) was incubated with vehicle, AB8939 (1-10 µM), paclitaxel (10 µM), or nocodazole (10 µM) at 37°C. AB8939 inhibits polymerization dose-dependently, with 50% inhibition at 1-2 µM and complete inhibition at ≥5 µM. Data represent three independent experiments. Panel C: Effect on pre-formed microtubules. Tubulin was polymerized for 30 minutes before AB8939 (10 µM) addition (arrow), demonstrating rapid disruption of established microtubules. Mean ± SEM from three experiments. AB8939 is a potent microtubule-destabilizing agent that directly inhibits tubulin polymerization.

To assess whether AB8939 directly affects tubulin polymerization, we performed a cell-free in vitro assay using porcine brain tubulin (Tubulin polymerization assay cat. BK011P, Cytoskeleton, Inc., Denver, USA). As demonstrated in Figure 1B, AB8939 inhibits in vitro tubulin polymerization in a dose-dependent manner compared to the solvent (DMSO) control. Fifty percent inhibition of microtubule (MT) polymerization was obtained at concentrations between 1 and 2 µM AB8939, and 100% inhibition at concentrations of 5 µM and above, comparable to the effect of nocodazole used at 10 µM. The microtubule-stabilizing agent (MSA) paclitaxel showed an early and increased polymerization rate compared to the control DMSO sample, as expected. In addition, Figure 1C shows that AB8939 can readily disrupt already-formed microtubules.

Therefore, the inhibition of tubulin polymerization and MT disruption in vitro correlates with the disruption of the microtubule network observed in AB8939-treated cells using immunofluorescence analysis (Figure 1). Disruption of the MT network is a hallmark of MTAs. By perturbing MT dynamics, MTAs interfere with mitotic spindle formation, arrest cells in G2/M, and eventually promote apoptotic cell death.

### AB8939 Targets the Colchicine-Binding Site of Tubulin

To understand how AB8939 interferes with tubulin properties at the molecular level, we determined the structure of the tubulin–AB8939 complex. Crystals of T2R, an assembly consisting of two αβ-tubulin heterodimers and one stathmin-like domain of the RB3 protein, were soaked in an AB8939-containing solution. Data up to 2.55 Å resolution were collected at the SOLEIL synchrotron (Supplementary data, Table S2). An electron density signal in which AB8939 could be modeled was identified at the colchicine-binding site (Figure 2). The electron density map enabled us to determine unambiguously the position and orientation of AB8939 (Figure 2B).

**Figure 2.**
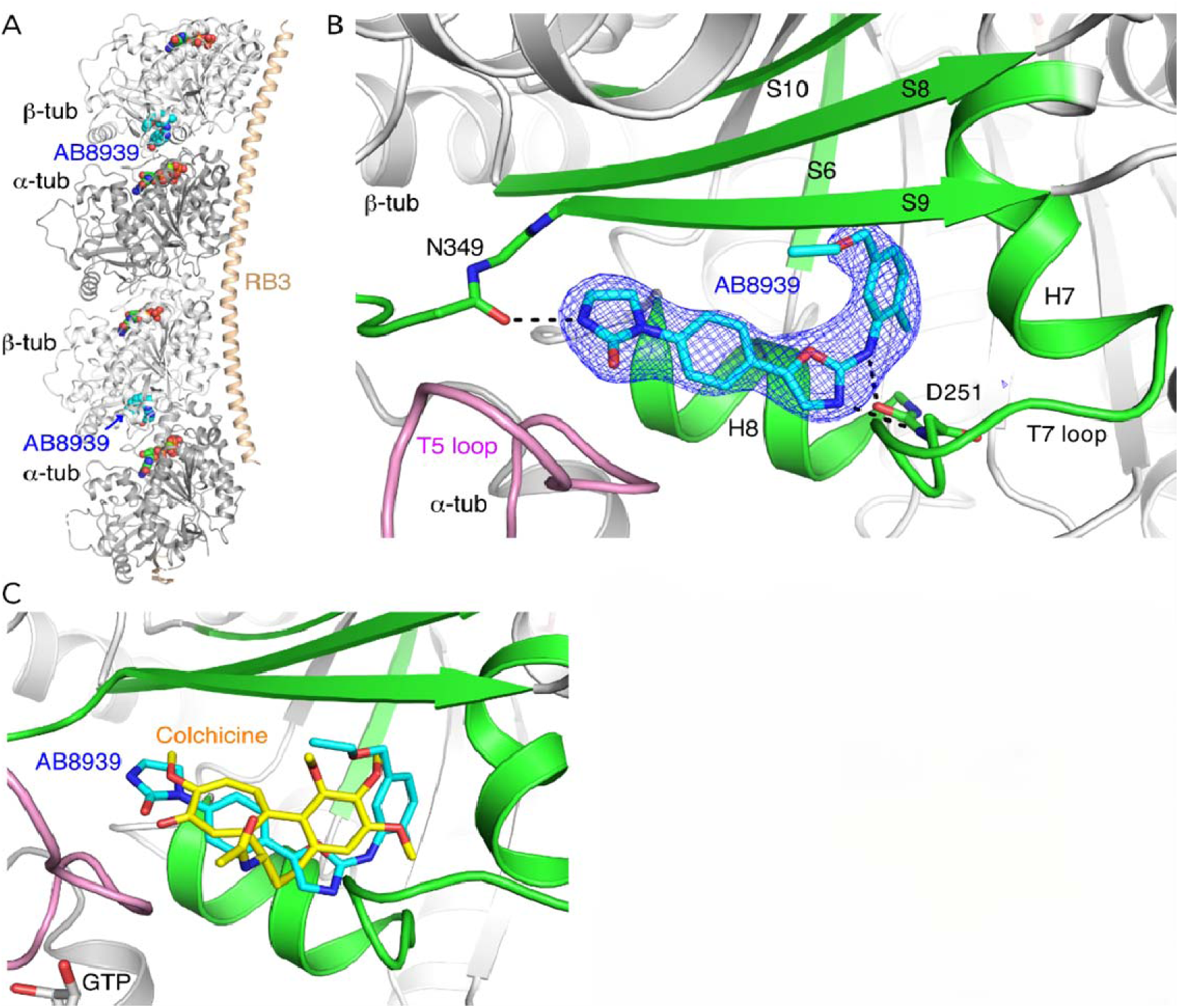
A B8939 Targets the Colchicine-Binding Site. Panel A: Crystal structure (https://www.ebi.ac.uk/pdbe/entry/pdb/8cgz) of T2R complex (two αβ-tubulin heterodimers and RB3 stathmin domain) bound to AB8939 (cyan). AB8939 binds at the colchicine site between α- and β-tubulin. Structure solved at 2.55 Å resolution (R_work/R_free = 19.8%/23.4%). Panel B: Detailed view of AB8939 binding site with F_obs-F_calc omit map (3σ, blue mesh). Key interactions include hydrogen bonds with β-tubulin Asp251 and Asn349, plus extensive van der Waals contacts. Panel C: Structural superposition of AB8939-bound (cyan/green) and colchicine-bound tubulin (PDB 5EYP, yellow/gray). Both occupy the same binding pocket with significant overlap. RMSD = 0.42 Å for binding site Cα atoms.

Similar to other colchicine-binding site inhibitors (CBSIs), such as colchicine (Figure 2C, (12)) and VERU-111 (data not shown, (13)), AB8939 binds to β-tubulin at the interface with the α subunit. The AB8939-binding site is formed mainly by residues of strands S8 and S9, loop T7, and helices H7 and H8 of β-tubulin, and to a lesser extent by loop T5 of α-tubulin and by strands S6 and S10 of the β subunit. In addition to van der Waals interactions, hydrogen bonds are engaged between AB8939 and the main chain atoms of β-tubulin residues Asp251 and Asn349 (Figure 2B).

These structures confirm AB8939 inhibits tubulin polymerization by binding the colchicine site, preventing conformational changes required for microtubule assembly.

### AB8939 Blocks Cell Cycle in G2/M and Promotes Apoptosis

To investigate whether AB8939 interferes with cell cycle progression, we performed cell cycle analysis. The effect of AB8939 on the cell cycle was evaluated using HCT116 cells (a human colorectal tumor cell line sensitive to AB8939, Table 1), treated with various concentrations of AB8939 for 24 hours. After permeabilization, the cells were analyzed for DNA content using propidium iodide staining. As seen in Figure 3A, AB8939 produced strong mitotic arrest in HCT116 cells after 24 hours of exposure, starting at concentrations as low as 3 nM, with more than 90% of treated cells arrested in G2/M phase at 10 nM and higher concentrations. The activity of AB8939 at 10 nM and 100 nM is identical to that of vincristine (Figure 3B).

**Figure 3.**
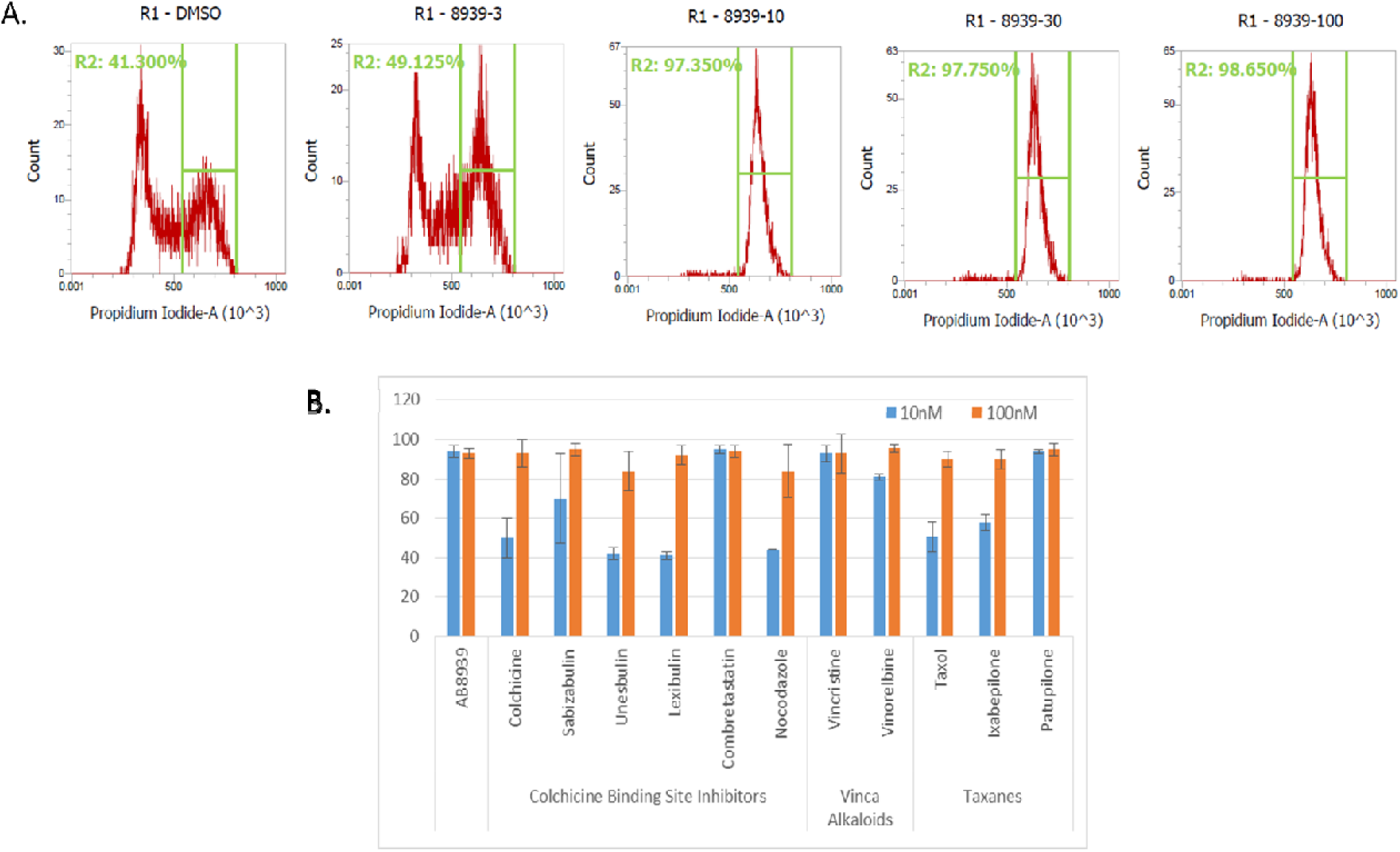
AB8939 induces a G2/M arrest. Panel A: Cell cycle profiles of HCT116 colorectal carcinoma cells treated for 24 hours with vehicle, AB8939 (3-100 nM), or vincristine (100 nM). Cells were stained with propidium iodide and analyzed by flow cytometry. AB8939 induces dose-dependent G2/M arrest, with >90% arrest at ≥10 nM, comparable to vincristine. Data represent three experiments. Panel B: Quantitative comparison of G2/M arrest by AB8939 versus other microtubule-targeting agents (vincristine, vinorelbine, ixabepilone, unesbulin, sabizabulin) at 10 nM and 100 nM. Mean ± SEM from three experiments. AB8939 demonstrates comparable or superior potency. One-way ANOVA with Dunnett’s test (*p < 0.05, **p < 0.01, ***p < 0.001). G2/M arrest was reversible at 10 nM but irreversible at ≥100 nM after 24-hour washout, indicating concentration-dependent cytostatic versus cytotoxic effects.

The comparison with other known MTAs shows that AB8939 is equivalent to or even more potent than most of these agents in inducing G2/M block. Tested microtubule-targeting agents include the approved drugs vincristine, vinorelbine, and ixabepilone, and others currently being evaluated in clinical development such as unesbulin (PTC596) and sabizabulin (VERU-111). The G2/M block induced by AB8939 is completely reversible at 10 nM following washout of 24 hours and irreversible at higher concentrations (100 nM and 1 µM) (data not shown).

The impact of AB8939 on G2/M phase cell cycle arrest and subsequent induction of apoptosis was further assessed using the Ara-C-resistant MOLM-14 cell line (14). MOLM-14 was originally derived from the peripheral blood of a 20-year-old male patient diagnosed with relapsed acute myeloid leukemia (AML), FAB subtype M5a, in 1995, following a prior history of myelodysplastic syndrome (MDS; refractory anemia with excess blasts, RAEB). This cell line harbors an internal tandem duplication in FLT3 and a ΔExon8 mutation in the CBL gene.

Consistent with the activity of most microtubule-targeting agents, the G2/M arrest leads to apoptosis, which can be distinguished into early stages—characterized by annexin V binding—and late stages, marked by propidium iodide uptake that reflects loss of membrane integrity and irreversible cellular damage.

Figure 4 presents representative flow cytometry data illustrating the apoptotic response of MOLM-14 tumor cells to AB8939 treatment. In particular, Figure 4A highlights the pronounced proapoptotic activity of AB8939, where a brief 3-hour exposure followed by washout and 24-hour analysis resulted in 12.3% late apoptosis. These findings demonstrate that even short-term treatment is sufficient to eliminate a substantial fraction of tumor cells.

**Figure 4.**
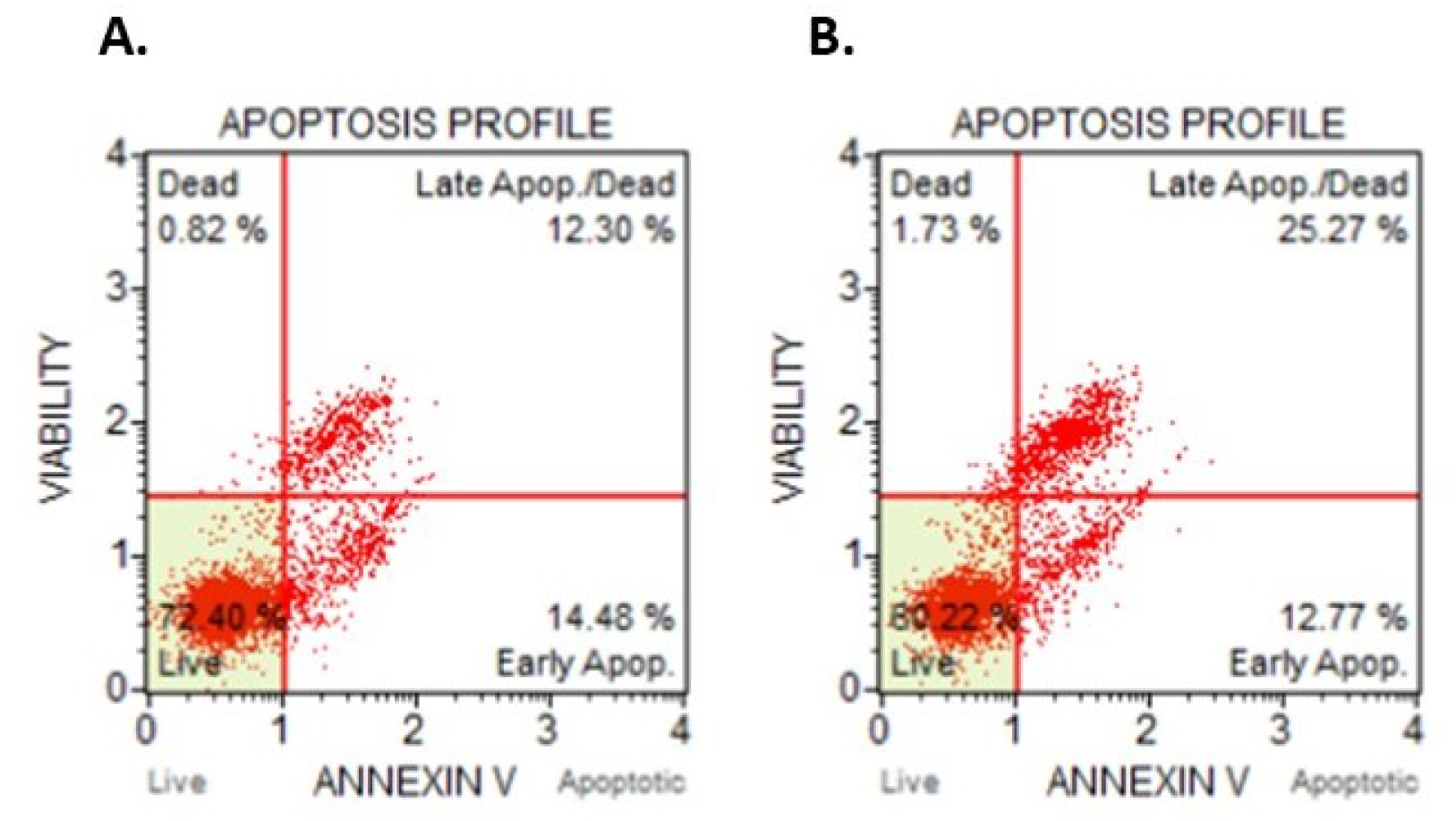
AB8939 Induces Apoptosis in AML MOLM-14 Cells. MOLM-14 cells (AML FAB M5a with FLT3-ITD and CBL mutations) were analyzed by Annexin V-FITC/PI staining. Quadrants show viable (Annexin V⁻/PI⁻), early apoptotic (Annexin V⁺/PI⁻), late apoptotic (Annexin V⁺/PI⁺), and necrotic (Annexin V⁻/PI⁺) populations. Panel A: 3-hour pulse treatment. Cells treated with 50 nM AB8939 for 3 hours, washed, and cultured 21 hours in drug-free medium. 12.3% underwent late apoptosis by 24 hours, demonstrating potent proapoptotic activity after brief exposure. Panel B: Continuous treatment for 24 hours with 50 nM AB8939 shows marked increase in both early and late apoptosis versus pulse treatment. 89.4% apoptotic cells with continuous exposure versus 45.7% with pulse treatment. Data from three experiments. Paired t-test: ***p < 0.001. AB8939 effectively induces apoptosis in chemo-resistant AML cells even after brief exposure, with enhanced effect during continuous treatment.

In a separate series of experiments, the proportion of cells undergoing G2/M arrest and subsequent apoptosis was quantified following treatment with AB8939 and other microtubule-targeting agents (MTAs). As summarized in Table 4, AB8939 induced a G2/M cell cycle block with efficacy comparable to that of established MTAs, while demonstrating clear superiority over ABT-751—a novel sulfonamide antimitotic targeting the colchicine-binding site of β-tubulin—and vincristine, a widely used G2/M blocker in the treatment of acute leukemias, including both acute lymphoblastic leukemia and acute myeloid leukemia.

**Table 4:** Effects of AB8939 and known microtubule targeting agents on G2/M phase cell cycle arrest and apoptosis in MOLM14 cells in vitro. Cells were treated for 24 hours.

| Drug | Concentration (nM) | Cells in G2/M (%) | Apoptosis (%) |  |
| --- | --- | --- | --- | --- |
|  |  |  | Early | Late/Dead |
| DMSO (control) | 0 | 24 | 7 | 3 |
| AB8939 | 10 | 43 | 31 | 15 |
| ABT-751 | 10 | 24 | 5 | 5 |
| Combretastatin A-4 | 10 | 43 | 30 | 17 |
| Lexibulin | 10 | 34 | 19 | 12 |
| Nocodazole | 10 | 25 | 4 | 1 |
| Ixabepilone | 10 | 56 | 21 | 7 |
| Patupilone | 10 | 55 | 27 | 17 |
| Plinabulin | 10 | 58 | 26 | 10 |
| Vincristine | 10 | 26 | 4 | 2 |
| Vinorelbine | 10 | 48 | 28 | 14 |

As shown in Table 4, AB8939 induces both early and late apoptosis at levels markedly higher than those observed with ABT-751, vincristine, or lexibulin—a novel microtubule polymerization inhibitor currently in phase 2 clinical trials. Notably, all cell lines exposed to AB8939 displayed G2/M arrest after 24 hours of treatment, indicating a consistent mechanism of action. In hematopoietic cell lines, this G2/M block rapidly progresses to apoptosis and cell death, whereas in non-hematopoietic cell lines, extended exposure of approximately 5–6 days is required (Supplementary data, Table S1).

### AB8939 Overcomes P-Glycoprotein (P-gp)-Mediated Drug Resistance

P-glycoprotein (P-gp), also known as ATP-binding cassette subfamily B member 1 (ABCB1), is a key mediator of multidrug resistance (MDR) in cancer. This membrane-associated transporter actively expels a broad spectrum of chemotherapeutic agents from cells in an ATP-dependent manner (15). Its overexpression, frequently observed in various malignancies including leukemias, is strongly associated with poor prognosis and reduced treatment efficacy. Because P-gp transports multiple frontline agents—such as doxorubicin, vincristine, and paclitaxel—its activity poses a major barrier to effective chemotherapy. We therefore examined whether AB8939 could overcome P-gp–mediated resistance in tumor cells.

To directly evaluate whether AB8939 is a P-gp substrate, we employed the P-gp-Glo™ assay (Promega), which measures ATPase activity of recombinant human P-gp. Substrates typically stimulate ATPase activity, whereas poor substrates or non-substrates show minimal effect. Doxorubicin, a well-characterized P-gp substrate, served as the positive control, while combretastatin A4 (CA-4), a poor substrate (16), was used as the negative control. As shown in Figure 5 (left panel), AB8939 behaved as a poor P-gp substrate, comparable to CA-4, in contrast to the strong ATPase stimulation observed with doxorubicin.

**Figure 5.**
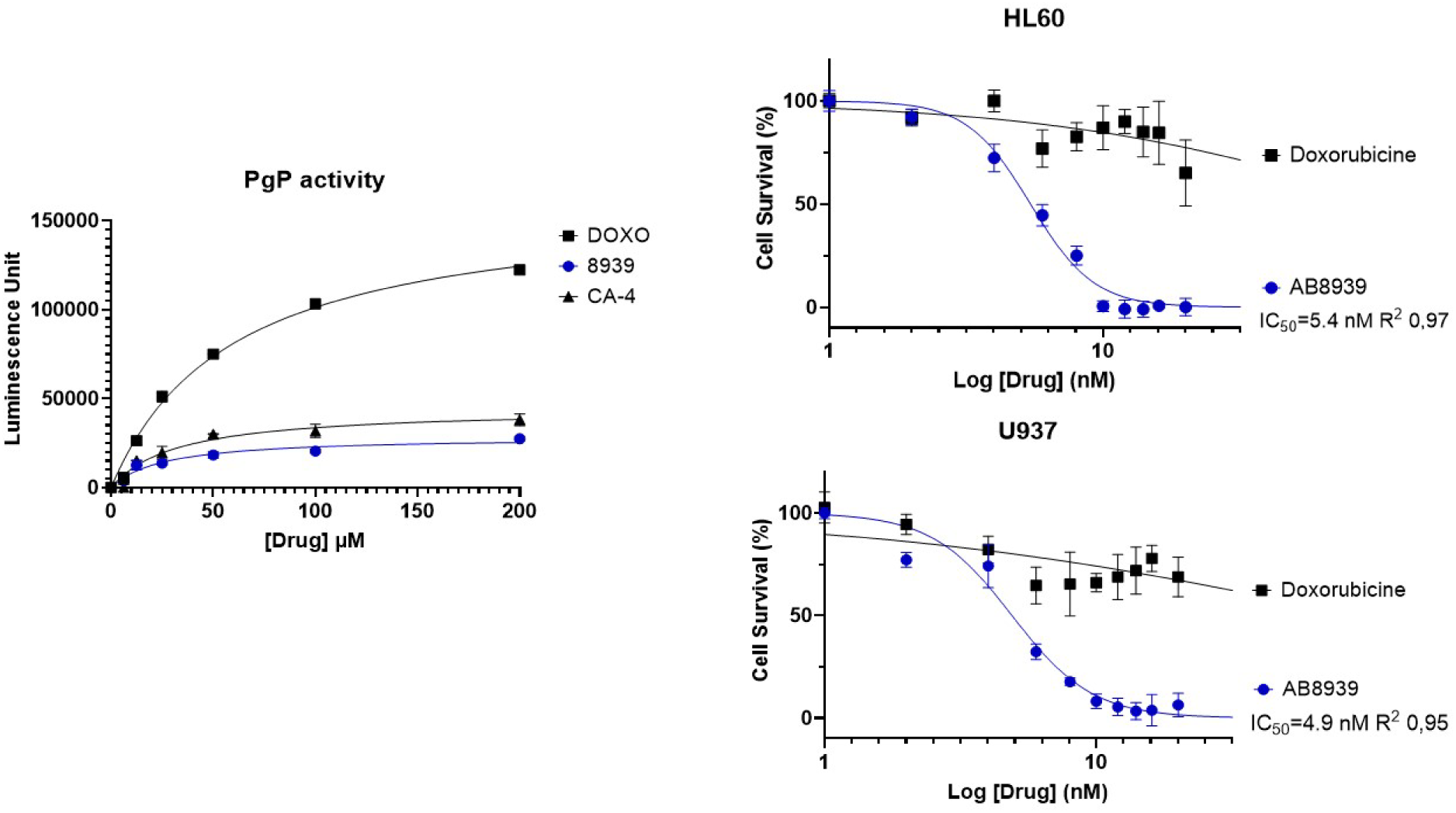
AB8939 Is Not a Substrate of P-Glycoprotein and Overcomes Multidrug Resistance. Panel A. P-glycoprotein (P-gp) ATPase activity measured with the P-gp-Glo™ Assay (Promega). Compounds (1–200 µM) were tested for effects on recombinant human P-gp (ABCB1). Luminescence (RFU) reflects ATPase modulation. Doxorubicin (black squares) strongly stimulated ATPase activity, peaking at 50–100 µM. AB8939 (blue circles) showed minimal stimulation, comparable to combretastatin A-4 (CA-4, black triangles), indicating poor recognition by P-gp. Data are mean ± SEM from three independent experiments (duplicate). Panel B. Antiproliferative activity of AB8939 versus doxorubicin in P-gp-positive AML cell lines. HL-60 and U-937 cells were treated for 72 h with increasing drug concentrations, and viability was assessed by CellTiter-Blue. AB8939 (blue circles) inhibited proliferation with IC₅₀ ≈ 5 nM in both lines, whereas doxorubicin (black squares) showed IC₅₀ >1000 nM, consistent with P-gp-mediated resistance. Data are mean ± SEM from three independent experiments (triplicate). IC₅₀ values were determined by nonlinear regression using GraphPad Prism 10.6.1.

The ability of AB8939 to overcome P-glycoprotein–mediated resistance was further evaluated in HL60 and U937 cell lines, both of which express P-gp (Figure 5, right panel). AB8939 potently inhibited proliferation in both models, with IC₅₀ values of approximately 5 nM. In contrast, at comparable concentrations, both cell lines exhibited strong resistance to doxorubicin.

To extend these findings, we compared the antiproliferative activity of AB8939 in the P-gp–negative MES-SA sarcoma cell line and its P-gp–positive variant, MES-SA/Mx2 (17). As summarized in Table 5, AB8939, like other agents not subject to P-gp efflux, showed equivalent activity in both strains. By contrast, drugs that are efficient P-gp substrates demonstrated markedly reduced efficacy in the resistant MES-SA/Mx2 cells. These in vitro data clearly show that AB8939 is a poor substrate of P-gp and overcomes multidrug resistance mediated by P-gp.

**Table 5.** Comparative anti-proliferative activity of AB8939 vs drugs known to be subtrate of P-gp and drugs known to be non-P-gp substrate on parental MES-SA and drug-resistant MES-SA/Mx2 (mdr1 positice) cell lines.

| DRUGS | MES-SA | MES-SA/Mx2 | RI |
| --- | --- | --- | --- |
| AB8939 | 5.7 +/- 5 | 5.2 +/- 4.7 | 1 |
| Colchicine | 8.6 +/- 1.2 | 11 +/- 1.5 | 1 |
| Combretastatine A4 | 1.6 +/- 0.5 | 1.5 +/- 0.7 | 1 |
| Paclitaxel | 8.0 +/- 5 | 3100 +/- 2690 | 400 |
| Vincristine | 7.9 +/- 6 | 3566 +/- 1808 | 450 |
| Doxorubicin | 71 +/- 56 | 4440 +/- 792 | 63 |

The IC<sub>50</sub> values expressed in nM are presented as a means of independent experiment ± standard deviation. RI=resistance index is calculated by dividing IC<sub>50</sub> of compound in resistant cell line by the IC<sub>50</sub> of corresponding parental cell line.

Beyond drug efflux, alternative mechanisms of resistance to microtubule-targeting agents (MTAs) have been linked to elevated β3-tubulin expression, which is also associated with aggressive disease phenotypes (18,19). To assess whether AB8939 retains activity in this context, we examined β3-tubulin expression across a panel of 33 human tumor cell lines by Western blot. High β3-tubulin levels were observed in most epithelial-derived lines, particularly those of lung, pancreatic, ovarian, and glioblastoma origin, whereas hematopoietic cell lines were largely negative, with the exception of the T-cell lymphoma line Karpas-299 (Supplementary data, Figure S2 and Table S1). Importantly, AB8939 potently inhibited proliferation in all 33 cell lines, with IC₅₀ values in the nanomolar range (Table S1). These findings indicate that AB8939 remains active in β3-tubulin–overexpressing tumors and, taken together with prior results, demonstrate its ability to overcome resistance mechanisms mediated both by drug efflux pumps and β3-tubulin expression.

### Ex Vivo Antiproliferative Activity of AB8939 on AML Blasts Isolated from Patients

Given the pronounced activity of AB8939 against hematopoietic tumor cell lines in vitro proliferation assays, we next evaluated its cytotoxic potential in primary blasts from AML patients. Peripheral blood (PB) and/or bone marrow (BM) samples were collected at diagnosis or relapse, following informed consent (HematoBio, NCT05602168; CeGAL, NCT02619071; Department of Hematology, Institut Paoli-Calmettes, France). After purification, mononuclear cells from PB or BM were exposed for 48 hours to increasing concentrations of AB8939, vincristine, 5-azacytidine (Vidaza), or cytarabine (Ara-C), and cell viability was subsequently assessed (Supplementary Table S3).

Figure 6 demonstrates that most AML blasts are resistant to vincristine (>2 µM, relative IC₅₀ = 100). Notably, among these vincristine-resistant samples, 45% of naïve AML blasts (Figure 6A, 6C) and over 33% of refractory/relapsed AML blasts (Figure 6B, 6D) remained sensitive to AB8939. All refractory/resistant AML blasts with IC₅₀ values >5 µM were also resistant to vincristine. Overall, AB8939 exhibited potent cytotoxic activity against patient-derived AML blasts, with IC₅₀ values ranging from 10 nM to 2 µM, including samples resistant to vincristine and the standard-of-care agent cytarabine (Ara-C). Importantly, strong activity was also observed in blasts from poorly differentiated AML-M1 subtypes, AML secondary to MDS, and monocytic AML (FAB-M5), across both naïve and relapsed/refractory cases (Figure 6), many of which harbored complex karyotypes (Supplementary data, Table S3). Strikingly, all samples carrying adverse-prognosis TP53 or ASXL1 mutations, or MECOM rearrangements, were sensitive to AB8939 compared with standard therapies (Ara-C and 5-azacytidine), suggesting that neither high-risk mutations nor unfavorable karyotypes diminish AB8939 activity.

**Figure 6.**
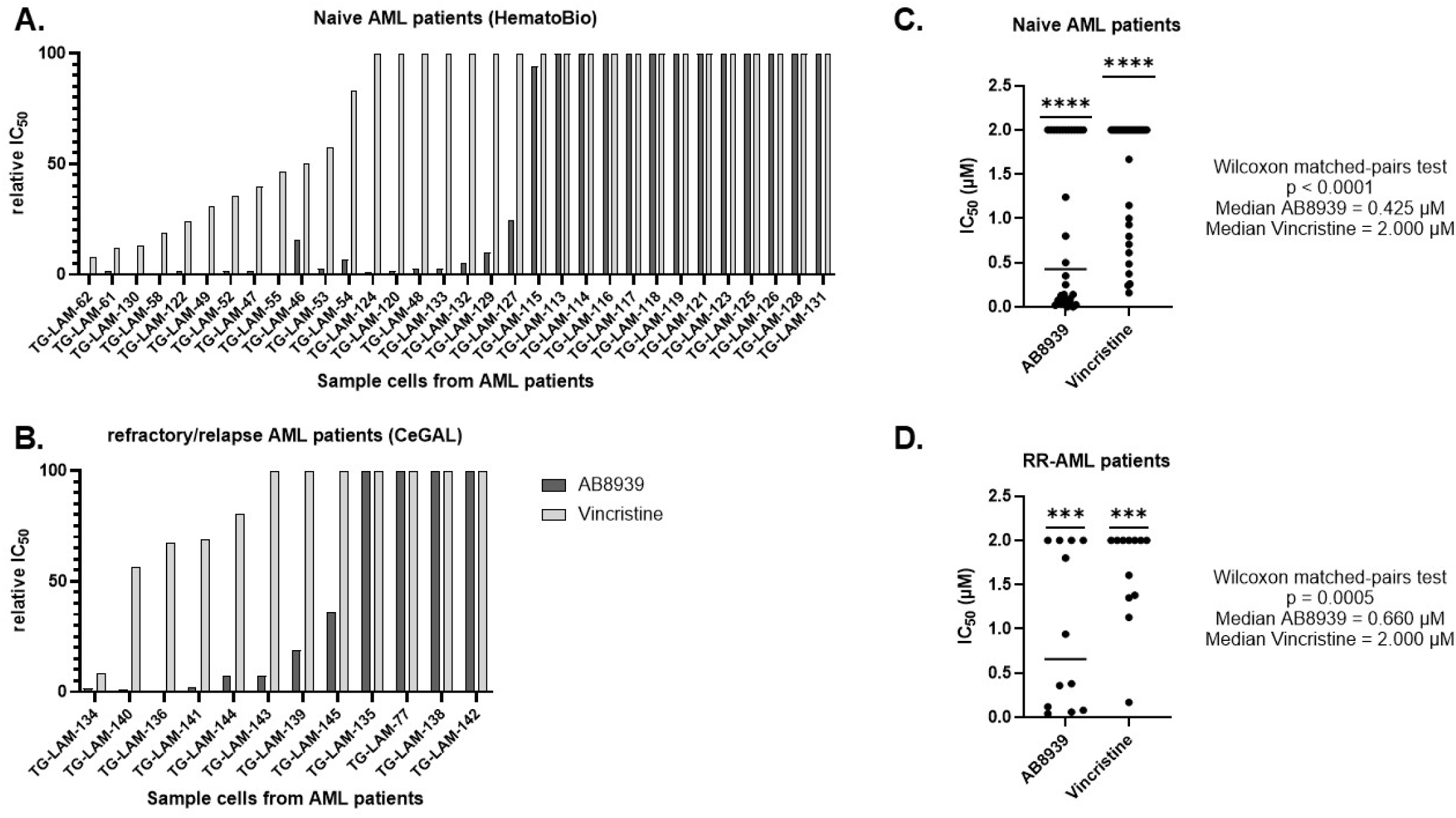
AB8939 Exhibits Potent Cytotoxicity Against Primary AML Blasts, Including Vincristine-Resistant and Refractory/Relapsed Cases. Ex vivo cytotoxicity of AB8939 and vincristine was assessed in primary acute myeloid leukemia (AML) blasts isolated from patient peripheral blood (PB) or bone marrow (BM) samples obtained at diagnosis (newly diagnosed, ND-AML) or at relapse/progression (refractory/relapsed, RR-AML) with informed consent (HematoBio NCT05602168; CeGAL NCT02619071, Institut Paoli-Calmettes, Marseille, France). Following Ficoll density gradient purification, mononuclear cells were treated for 48 h with serial dilutions of AB8939 or vincristine, and viability was measured using the CellTiter-Glo assay (Promega). Panel A: Cytotoxicity profiles of ND-AML patient samples (n = 32). Each row represents an individual patient; columns show relative IC₅₀ values for AB8939 (dark grey) or vincristine (light grey), calculated as ((IC_50_ of the drug µM/Max drug concentration in the assay (AB8939 5 µM, Vincristine 2 µM)*100). Many vincristine-resistant samples (relative IC₅₀ = 100, i.e. >2 μM) remained sensitive to AB8939 (IC₅₀ < 500 nM), indicating AB8939 can overcome vincristine resistance. Panel B: Cytotoxicity profiles of RR-AML patient samples (n = 12). Despite broad resistance to vincristine and standard-of-care agents (see Supplementary Table S1), RR-AML blasts displayed marked sensitivity to AB8939. Panel C: Comparative IC₅₀ values in ND-AML samples (n = 32). Each dot represents an individual patient. Median IC₅₀ values: AB8939 = 0.425 μM vs. vincristine = 2.000 μM. Wilcoxon matched-pairs signed-rank test confirmed significantly greater potency of AB8939 (p < 0.0001; sum of signed ranks = 528.0). Panel D: Comparative IC₅₀ values in RR-AML samples (n = 12). Median IC₅₀ values: AB8939 = 0.660 μM vs. vincristine = 2.000 μM. Wilcoxon matched-pairs signed-rank test again demonstrated superior potency of AB8939 (p = 0.0005; sum of signed ranks = 78.0). Overall, AB8939 consistently outperformed vincristine across both naïve and refractory/relapsed AML patient samples, including highly chemoresistant cases.

### AB8939 Has Antitumor Activity In Vivo in a Preclinical Model of Human AML

Building on the encouraging in vitro and ex vivo activity of AB8939 against AML cell lines and primary blasts, we next investigated its therapeutic potential in vivo using a chemotherapy-resistant mouse model (MOLM-14) engineered to express luciferase (MOLM-14-Luc). Consistent with our earlier findings (Table 1 and Figure 4), AB8939 displayed potent cytotoxicity against MOLM-14-Luc cells in vitro, with an IC₅₀ of 17 nM, whereas the same cells were resistant to the standard-of-care agent cytarabine (Ara-C; IC_50_ = 200 nM, data not shown).

The in vivo study design is outlined in Figure S3 (Supplementary Data). Briefly, on day 0 (D0), 0.2 × 10LJ MOLM-14-Luc cells were injected intravenously into immunodeficient NSG mice. On day 1 (D1), animals were randomized into seven groups of four. Group 1 received vehicle control (20% hydroxypropyl β-cyclodextrin in water) administered subcutaneously (SC) for six consecutive days per week (6 on/1 off). The remaining groups were treated with AB8939 as a single agent, delivered SC at doses of 1, 3, or 6 mg/kg, following either a 3 on/4 off or 6 on/1 off schedule. Animals in the 3 mg/kg cohort were sacrificed on day 33 for ethical reasons, following the decision to continue treatment only at the 6 mg/kg dose. The 6 mg/kg (3 on/4 off) cohort was discontinued on day 37 at the end of the sixth cycle because of disease progression, whereas the 6 mg/kg (6 on/1 off) cohort was maintained until day 53.

Leukemia progression was monitored weekly by bioluminescence imaging (Figure 7A). By day 22 post-injection, imaging revealed a clear dose-dependent antitumor effect of AB8939 in the MOLM-14-Luc AML model (Figure 7B, 7C). Kaplan-Meier survival analysis further demonstrated that no survival benefit was observed in the 1 mg/kg cohorts compared with vehicle, with both groups showing a median survival of ∼29 days (Figure 7D). In contrast, both 6 mg/kg cohorts exhibited significant survival advantages, with median survival extended to 40.5 days (3 on/4 off) and 59 days (6 on/1 off). Notably, all mice in the 6 mg/kg, 6 on/1 off group remained alive at Day 56, indicating durable disease control.

**Figure 7.**
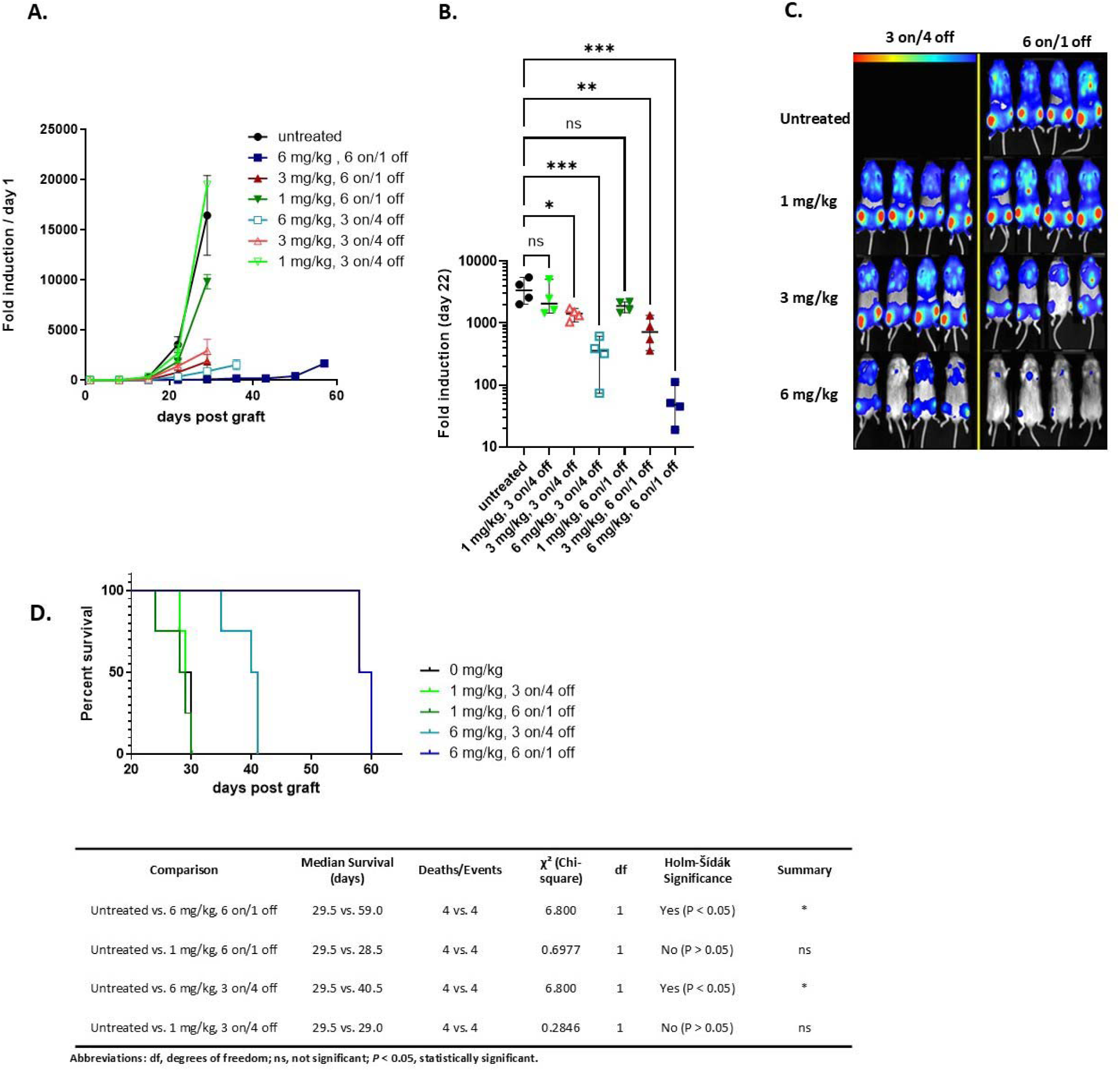
Antitumor activity of AB8939 in vivo in a preclinical AML model. Female NSG mice were intravenously engrafted with MOLM-14-Luc cells and randomized into treatment groups (n = 4). Mice received vehicle or AB8939 (1, 3, or 6 mg/kg) subcutaneously on either a 3 on/4 off or 6 on/1 off schedule. (A) Longitudinal bioluminescence imaging revealed dose- and schedule-dependent leukemia suppression, with maximal efficacy in the 6 mg/kg, 6 on/1 off group. (B) Quantification at Day 22 confirmed significant tumor burden reduction at 3 mg/kg and near-complete control at 6 mg/kg (one-way ANOVA with Dunnett’s test: P < 0.05, **P < 0.001 vs. vehicle). (C) Representative images at Day 22 illustrate dose-dependent disease control. (D) Kaplan-Meier survival analysis showed no benefit at 1 mg/kg (median ∼29 days), whereas both 6 mg/kg regimens significantly prolonged survival compared with vehicle (40.5 vs. 59 days; Log-rank P < 0.01). Holm-Šídák’s test confirmed significance for both 6 mg/kg groups (χ² = 6.800, df = 1; P < 0.05), with the 6 on/1 off schedule yielding the most durable response (100% survival at Day 56).

Log-rank (Mantel-Cox) testing confirmed significant survival benefits for both 6 mg/kg regimens compared with vehicle (*P* < 0.01), with the 6 on/1 off schedule producing the most pronounced effect (*P* = 0.003 vs. vehicle; *P* = 0.01 vs. 3 on/4 off). To further validate these findings, Holm-Šídák’s multiple comparisons test was applied. Compared with untreated controls (median survival: 29.5 days), AB8939 at 6 mg/kg significantly prolonged survival under both dosing schedules: 6 on/1 off (median: 59.0 days; χ² = 6.800, df = 1; *P* < 0.05) and 3 on/4 off (median: 40.5 days; χ² = 6.800, df = 1; *P* < 0.05). In contrast, no significant benefit was observed at 1 mg/kg under either schedule (median: 28.5–29.0 days; *P* > 0.05). Across all dosing regimens, AB8939 was well tolerated when administered subcutaneously, with no detectable impact on body weight (Supplementary Data, Figure S4).

In summary, these findings demonstrate that single-agent AB8939 effectively delays disease progression and significantly prolongs survival in an Ara-C–resistant human AML xenograft model. Given that Ara-C–based combinations remain the backbone of AML induction therapy, these results provide preliminary in vivo proof of concept supporting AB8939 as a promising candidate for the treatment of refractory AML.

### AB8939 in Refractory AML with MECOM Rearrangement: Efficacy as Single Agent and in Combination with Azacitidine or Venetoclax

Azacitidine is widely used in the treatment of myelodysplastic syndromes and has also demonstrated clinical benefit in acute myeloid leukemia (AML), particularly in patients ineligible for intensive chemotherapy (20). Clinical trials have shown that azacitidine can extend overall survival and improve quality of life in selected AML populations, especially older patients or those harboring genetic alterations that influence treatment response. However, its efficacy as a single agent remains limited, prompting the development of combination strategies to enhance therapeutic outcomes.

In this context, we evaluated the antitumor activity of AB8939 alone, azacitidine (Vidaza) alone, and their combination in the TG-LAM-75 patient-derived xenograft (PDX) model. TG-LAM-75 was established from a patient with AML-M1 carrying a MECOM rearrangement, a high-risk cytogenetic abnormality associated with poor prognosis and therapy resistance. The blasts are poorly differentiated and express CD33, a marker present on leukemic blasts in ∼85–90% of AML cases. The patient had previously failed both cytarabine/idarubicin induction and second-line azacitidine therapy, making this PDX a clinically relevant model of high-risk, treatment-refractory AML.

For in vivo studies, NSG mice were engrafted with 0.1 × 10LJ TG-LAM-75 cells (Day 0), and treatment was initiated on Day 17 upon blast detection. AB8939 was administered subcutaneously at 6 mg/kg (5 days/week for 4 weeks, n=9), azacitidine intraperitoneally at 2 mg/kg (daily for 13 days, n=9), and venetoclax orally at 100 mg/kg (daily for 7 days, n=9). Combination groups (n=9 each) received concurrent dosing according to the single-agent schedules. Vehicle controls (n=5) received subcutaneous injections on the AB8939 schedule. Blast burden was monitored in peripheral blood (Days 14, 31, 45) and in bone marrow and spleen at study termination (Day 46) by flow cytometry. Statistical analyses were performed using one-way ANOVA and Mann–Whitney tests.

### Efficacy of AB8939 with Azacitidine

We evaluated the effects of AB8939 and azacitidine (Vidaza), administered either as single agents or in combination, on the proportion of CD33⁺ cells in blood, bone marrow, and spleen. AB8939 monotherapy produced blast reductions that were comparable to or greater than those achieved with azacitidine in peripheral blood (Figure 8A), bone marrow (Figure 8B), and spleen (Figure 8C). After four weeks of treatment, both agents significantly decreased blast burden relative to vehicle controls. Notably, the combination of AB8939 with azacitidine resulted in near-complete disease clearance, with blast counts approaching zero across all compartments (Figure 8A–C). This combination regimen demonstrated significantly greater efficacy than either monotherapy, reducing circulating blasts to <5% and leaving only minimal residual disease in bone marrow and spleen.

**Figure 8.**
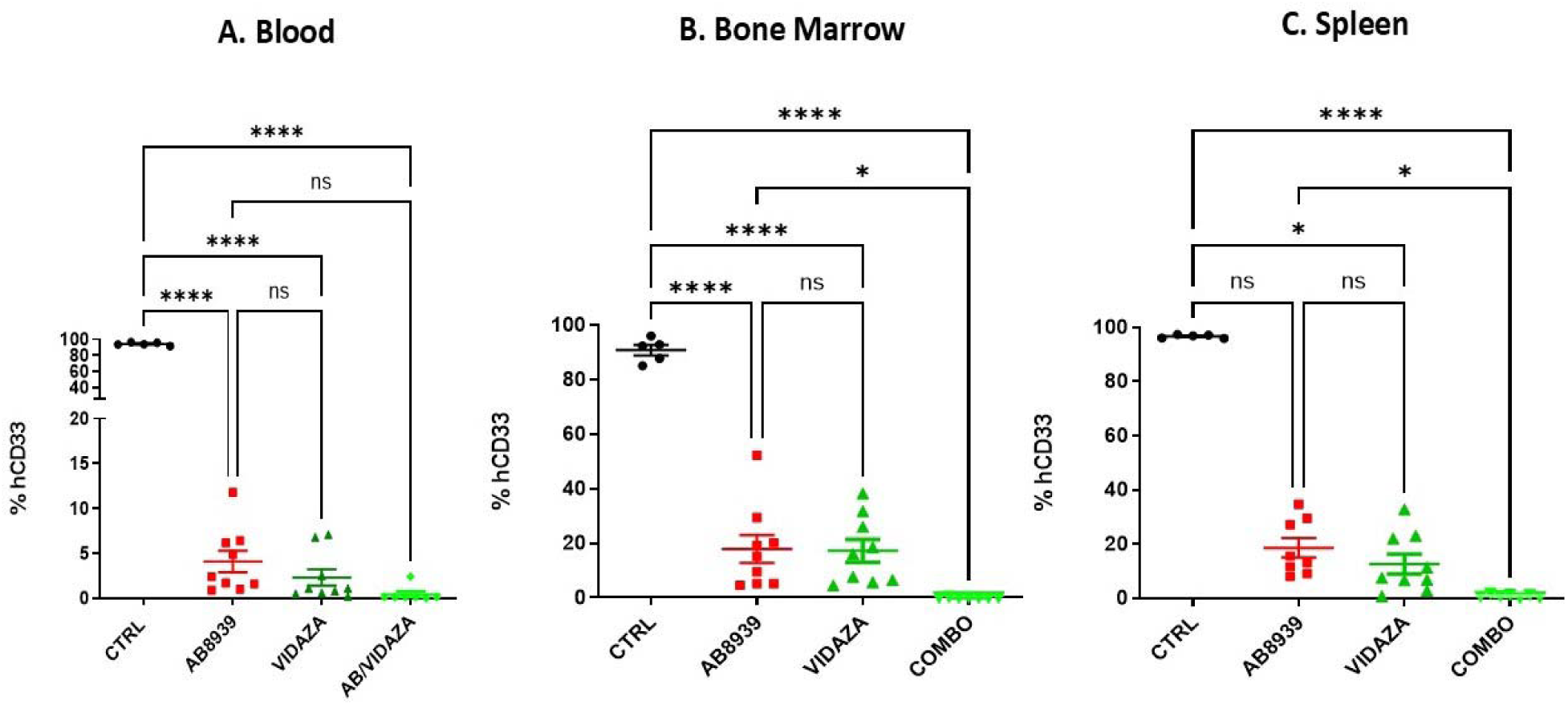
Anti-leukemic Efficacy of AB8939 with Azacitidine in the TG-LAM-75 PDX Model. Panel A: Percentage of human CD33^+^ blast cells in peripheral blood at Day 45 post-engraftment (28 days post-treatment initiation). Data show percentage of CD33^+^ cells relative to total leukocytes (100 × hCD33^+^/(hCD45^+^ + mCD45^+^)). The combination of AB8939 plus azacitidine reduced blast percentages to near-zero levels (<5%), significantly superior to either monotherapy. Panel B: Percentage of human CD33^+^ blast cells in bone marrow at Day 46 post-engraftment. Combination therapy achieved dramatic blast reduction in the bone marrow compartment, with minimal residual disease detected. Panel C: Percentage of human CD33^+^ blast cells in spleen at Day 46 post-engraftment. The combination treatment produced marked blast elimination in the spleen, demonstrating multi-compartment disease control. All measurements were performed by flow cytometry for human CD45^+^/CD33^+^ cells. Data are presented as individual values with mean ± SEM. Statistical analyses were performed using one-way ANOVA with post-hoc comparisons (*p < 0.05; **p < 0.01; ***p < 0.001). The combination therapy demonstrated significantly greater efficacy than either single agent across all tissue compartments.

### Safety Profile and Toxicity Assessment

Combination therapy is a standard strategy in leukemia treatment. For example, the cytarabine–doxorubicin regimen commonly used for AML induction is effective but often associated with severe toxicities, including myelosuppression, cardiotoxicity, and gastrointestinal complications. Thus, beyond assessing efficacy, it is critical to evaluate the safety of novel drug combinations in preclinical models.

Our results indicate that AB8939 displays a markedly improved safety profile compared with azacitidine. As shown in Figure 9A, azacitidine treatment caused profound depletion of murine CD45⁺ leukocytes (>80% reduction), consistent with the well-documented myelosuppressive effects of hypomethylating agents. In contrast, AB8939 preserved leukocyte counts at levels comparable to vehicle-treated controls, indicating an absence of haematotoxicity. Body weight monitoring (Figure 9B) further confirmed the favorable safety profile of AB8939, as treated mice maintained stable weights indistinguishable from controls. By comparison, azacitidine induced a significant decline in body weight, particularly between days 30 and 33 post-engraftment (13–16 days after treatment cessation). Importantly, the combination of AB8939 with azacitidine exhibited a toxicity profile similar to azacitidine alone, suggesting that AB8939 does not exacerbate the adverse effects of azacitidine.

**Figure 9.**
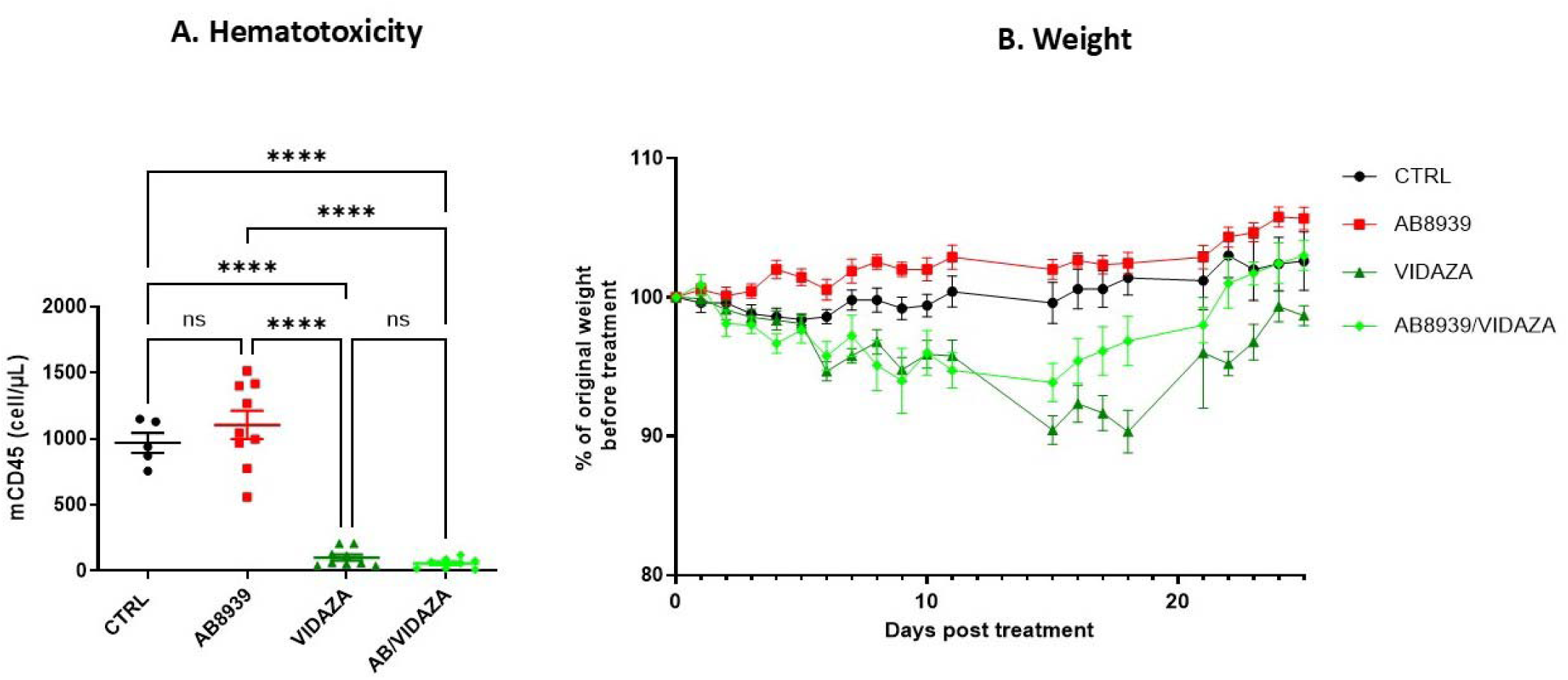
Safety Profile of AB8939 with Azacitidine. Tolerability was assessed throughout the study as described in Figure 7. Animals were monitored daily for clinical signs and body weight changes. Safety parameters were evaluated for vehicle control, AB8939 alone, azacitidine alone, and AB8939 plus azacitidine combination. Panel A: Murine CD45^+^ leukocyte quantification in peripheral blood (Day 31 post-engraftment, 14 days post-treatment initiation). Detection of murine CD45^+^ cells serves as a marker of myelosuppression and hematotoxicity. Azacitidine treatment induced profound depletion of murine leukocytes (>80% reduction), consistent with known myelosuppressive effects of hypomethylating agents. AB8939 treatment preserved murine leukocyte counts at levels comparable to vehicle control, indicating absence of hematotoxicity. The combination showed similar myelosuppression to azacitidine alone. Data are presented as absolute cell counts (mCD45^+^ cells/µL blood) with individual values and mean ± SEM (vehicle n=5; AB8939 n=9; azacitidine n=9; combination n=8). Statistical significance determined by one-way ANOVA (***p < 0.001 azacitidine vs. vehicle and AB8939). Panel B: Body weight monitoring over time. Mice were weighed at regular intervals from Day 0 (engraftment) through Day 46 (study termination). Body weights are expressed as percentage of initial body weight. Azacitidine treatment (both alone and in combination with AB8939) induced significant weight loss (∼10% body weight) beginning approximately 13 days post-treatment cessation (Day 30 onwards), indicating delayed toxicity. In contrast, AB8939 monotherapy showed no body weight changes compared to vehicle control throughout the 4-week treatment period and follow-up, demonstrating excellent tolerability at the 6 mg/kg dose. Data are presented as mean ± SEM (vehicle n=5; AB8939 n=9; azacitidine n=9; combination n=9). Note: In the combination arm, 2 of 9 mice treated with AB8939 plus azacitidine died during the study, suggesting the combination may carry additional toxicity risk beyond azacitidine alone, despite similar weight loss profiles. No deaths occurred in AB8939 monotherapy groups.

A possible explanation for the maintained efficacy despite azacitidine-associated toxicity is that azacitidine induces profound myelotoxicity (Figure 9A), whereas AB8939 selectively targets leukemic cells, thereby potentially allowing repopulation of the bone marrow by normal myeloid progenitors.

### Venetoclax Combination Studies

While venetoclax monotherapy exhibited minimal efficacy in this refractory model (Figure 10A–C), its combination with AB8939 showed a clear trend toward enhanced anti-leukemic activity. In both peripheral blood and bone marrow, the combination produced greater numerical reductions in blast burden compared with AB8939 alone (Figure 10A–B), and in the spleen, blast percentages were further decreased under combination treatment (Figure 10C). Although these differences did not reach statistical significance, the observed trends point to potential synergistic activity that merits further investigation.

**Figure 10.**
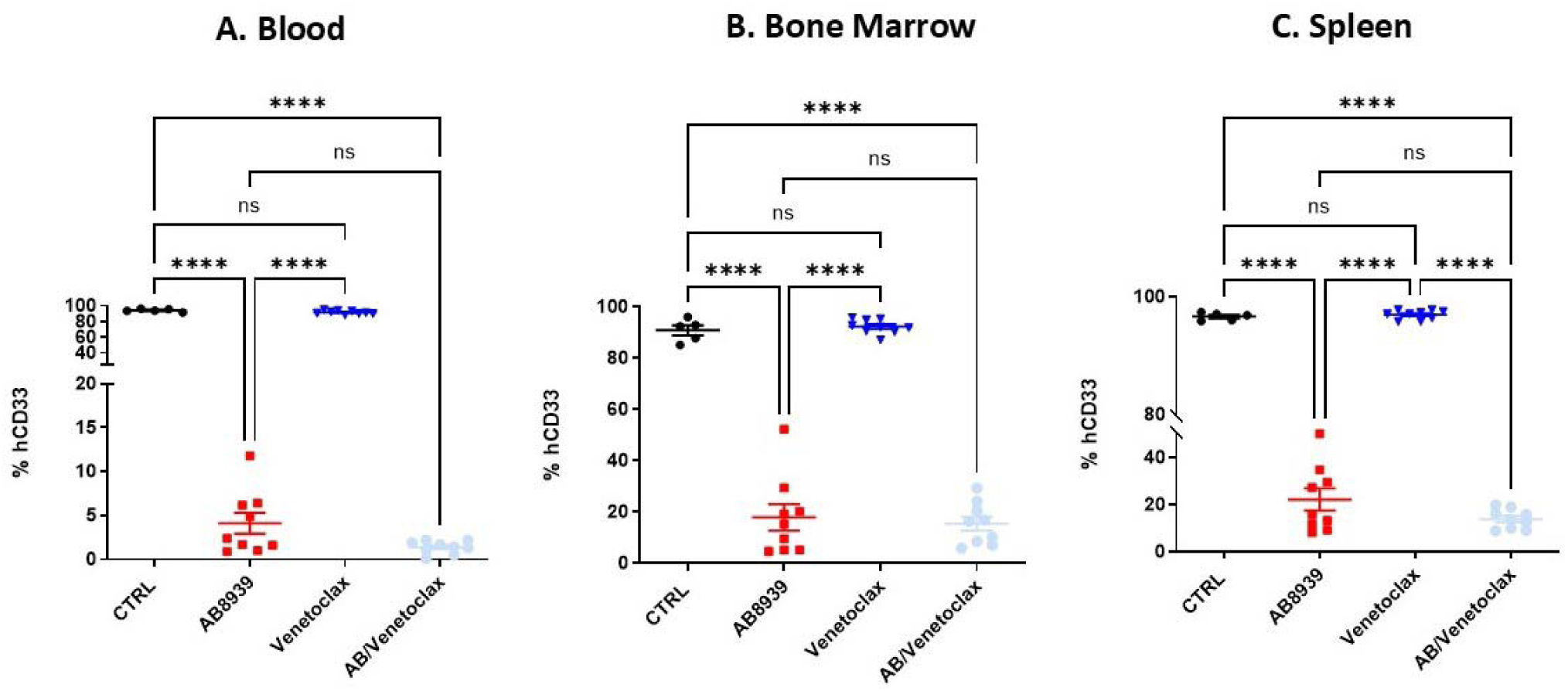
Efficacy of AB8939 with Venetoclax in TG-LAM-75 PDX model. NSG mice were engrafted with 0.1 × 10^6^ TG-LAM-75 PDX cells on Day 0. Treatment was initiated on Day 17. Animals received: vehicle control (n=5), AB8939 alone (6 mg/kg SC, 5 days/week for 4 weeks, n=9), venetoclax alone (100 mg/kg oral gavage daily for 7 consecutive days, n=9), or AB8939 plus venetoclax combination (n=9) using the same dosing schedules. Panel A: Percentage of human CD33^+^ blast cells in peripheral blood at Day 45 post-engraftment (28 days post-treatment initiation). Venetoclax monotherapy demonstrated minimal single-agent activity against the TG-LAM-75 PDX model. The combination of AB8939 plus venetoclax showed a trend toward greater blast reduction compared to AB8939 alone, though differences did not reach statistical significance. Panels B-C: Percentage of human CD33^+^ blast cells in bone marrow and spleen at Day 46 post-engraftment. Venetoclax alone had negligible effect. The AB8939/venetoclax combination showed numerical reduction in blast burden compared to AB8939 monotherapy, particularly evident in bone marrow, suggesting potential synergistic activity that warrants further investigation. All measurements were performed by flow cytometry. Data are presented as individual values with mean ± SEM. Statistical analyses performed using one-way ANOVA and Mann-Whitney t-test. While not statistically significant, the trends observed support further evaluation of the AB8939/venetoclax combination, and potentially a triplet regimen combining AB8939, azacitidine, and venetoclax.

Body weight monitoring (Supplementary Figure S9) confirmed that venetoclax alone, as well as in combination with AB8939, was well tolerated, with all treatment groups maintaining stable weights throughout the study. These findings demonstrate that the AB8939/venetoclax regimen does not introduce additional toxicity. Consistent with earlier results, AB8939 treatment also preserved murine leukocyte counts at levels comparable to vehicle controls, indicating an absence of hematotoxicity. Together, these data provide strong support for the continued development of AB8939.

Overall, the results suggest that AB8939 is a promising therapeutic candidate for azacitidine-refractory AML, including cases with high-risk genetic features such as MECOM rearrangements. The AB8939/azacitidine combination achieved potent disease control with a toxicity profile no greater than that of azacitidine alone. In addition, the favorable safety profile of AB8939 in combination with venetoclax highlights further therapeutic opportunities for patients with treatment-resistant AML.

### AB8939 Eradicates Blasts in Mouse Bone Marrow Bearing Aggressive AMKL Expressing ETO2/GLIS2 Fusion Protein

We further evaluated the in vivo antitumor activity of AB8939 using acute megakaryoblastic leukemia (AMKL) cells harboring the ETO2-GLIS2 translocation. This fusion oncogene, characteristic of aggressive AMKL and pediatric AML, drives leukemogenesis by enhancing self-renewal capacity and blocking differentiation, thereby maintaining cells in a primitive, undifferentiated state (10). The AMKL model employed in this study faithfully recapitulates the molecular features of human AMKL. Specifically, we used AMKL26 patient-derived xenograft (PDX) cells transduced with a luciferase/mCherry-expressing lentivirus, enabling longitudinal monitoring of leukemia progression by bioluminescence imaging.

Following confirmation of engraftment by imaging at two weeks post-injection, mice were randomized into treatment groups (non-blinded) and treated for three weeks with either vehicle or AB8939 (2 mg/kg i.v., 3 days ON/4 days OFF for 2 weeks, followed by 5 mg/kg, 3 days ON/4 days OFF for 1 week). Disease progression was monitored weekly by bioluminescence imaging (IVIS Spectrum). At seven weeks post-injection, bone marrow was harvested from femurs and tibiae for flow cytometry. Human leukemic burden was quantified as the percentage of mCherry-positive blasts among live cells (Sytox Blue exclusion, Invitrogen) in vehicle-versus AB8939-treated mice.

Figure 11A presents representative examples of the experimental results, with mice numbered 25–34 serving as controls and mice numbered 35–44 receiving AB8939 treatment according to the study design. Some heterogeneity in fluorescence intensity was observed, and the figure highlights the most illustrative outcomes. Figure 11B quantifies luminescence (photons/second) in control versus treated mice, showing a significant reduction in signal in the AB8939 group by the end of week 4. At weeks 5 and 6, however, the differences were no longer statistically significant, likely reflecting variability in luminescence measurements. Most notably, Figure 11C demonstrates that AB8939 completely eradicated AMKL blasts from the bone marrow in 75% of treated animals. Taken together, these findings underscore the potent efficacy of AB8939 against aggressive and refractory AML, including pediatric AMKL driven by the ETO2-GLIS2 fusion.

**Figure 11.**
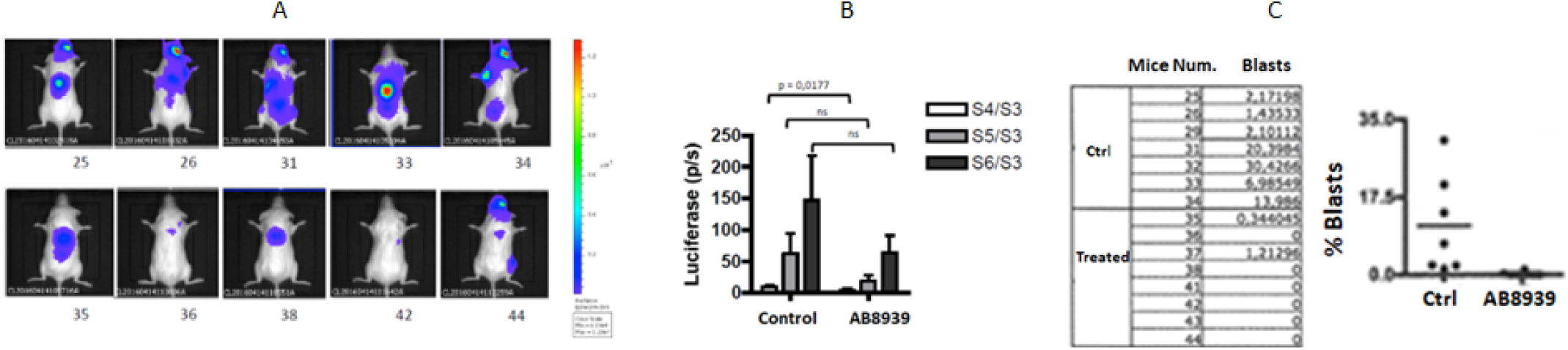
Anti-leukemic Efficacy of AB8939 with Azacitidine in the AMKL26 PDK model. NSG mice were engrafted with luciferase/mCherry-labeled AMKL26 PDX cells (Day 0), modeling aggressive pediatric AMKL. After confirming engraftment at Week 2, mice were randomized into vehicle (n=10, mice 25–34) or AB8939-treated (n=10, mice 35–44) groups. AB8939 was given IV over 3 weeks (2 mg/kg × 2 weeks, then 5 mg/kg × 1 week). Mice were monitored weekly by bioluminescence imaging (BLI) and euthanized at Week 7 for bone marrow analysis. Panel A: Representative BLI images at Week 4 show high leukemic burden in controls and reduced signal in AB8939-treated mice. Full dataset in Supplementary Fig. S9. Panel B: Longitudinal BLI quantification. AB8939 significantly reduced signal at Week 4 (***p < 0.001); differences at Weeks 5–6 were not significant due to inter-animal variability. Panel C: Flow cytometry at Week 7 revealed complete leukemic clearance (< 0.0001. See Supplementary Fig. S10.

### AB8939 Eradicates Leukemic Stem Cells in a Human PDX TG-LAM-36 AML Model

Relapse in acute myeloid leukemia (AML) remains a major clinical challenge, occurring in approximately 50% of patients who initially achieve remission. A key driver of relapse is minimal residual disease (MRD), often maintained by leukemic stem cells (LSCs) due to their intrinsic self-renewal capacity and resistance to conventional cytotoxic agents. To assess the impact of AB8939 on LSC persistence and leukemia recurrence, we employed the luciferase-expressing human AML PDX model TG-LAM-36, which is highly sensitive to AB8939 (IC_50_ = 40 nM) but fully resistant to cytarabine (5-Ara-C; IC₅₀ > 20 µM) in vitro.

NSG mice were engrafted with TG-LAM-36 cells and treated for four days with AB8939, 5-Ara-C, or their combination. Following treatment, human leukemic cells (hCD45⁺) were harvested and analyzed for LSC content using CD34⁺/CD38⁻/CD123⁺/JAM-C⁺ markers. These hCD45⁺ cells were then re-transplanted into naïve NSG mice at limiting dilutions (10,000; 2,000; 400; 80 cells), and leukemia recurrence was monitored over 81 days without additional therapy.

Flow cytometry revealed a significant reduction in total hCD45⁺ cells in the bone marrow of both AB8939- and Ara-C–treated mice (Figure 12A). Notably, AB8939 treatment resulted in a relative enrichment of CD34⁺/CD38⁻ LSCs within the residual hCD123⁺ population (Figure 12B), likely reflecting more efficient depletion of proliferating blasts compared with Ara-C.

**Figure 12.**
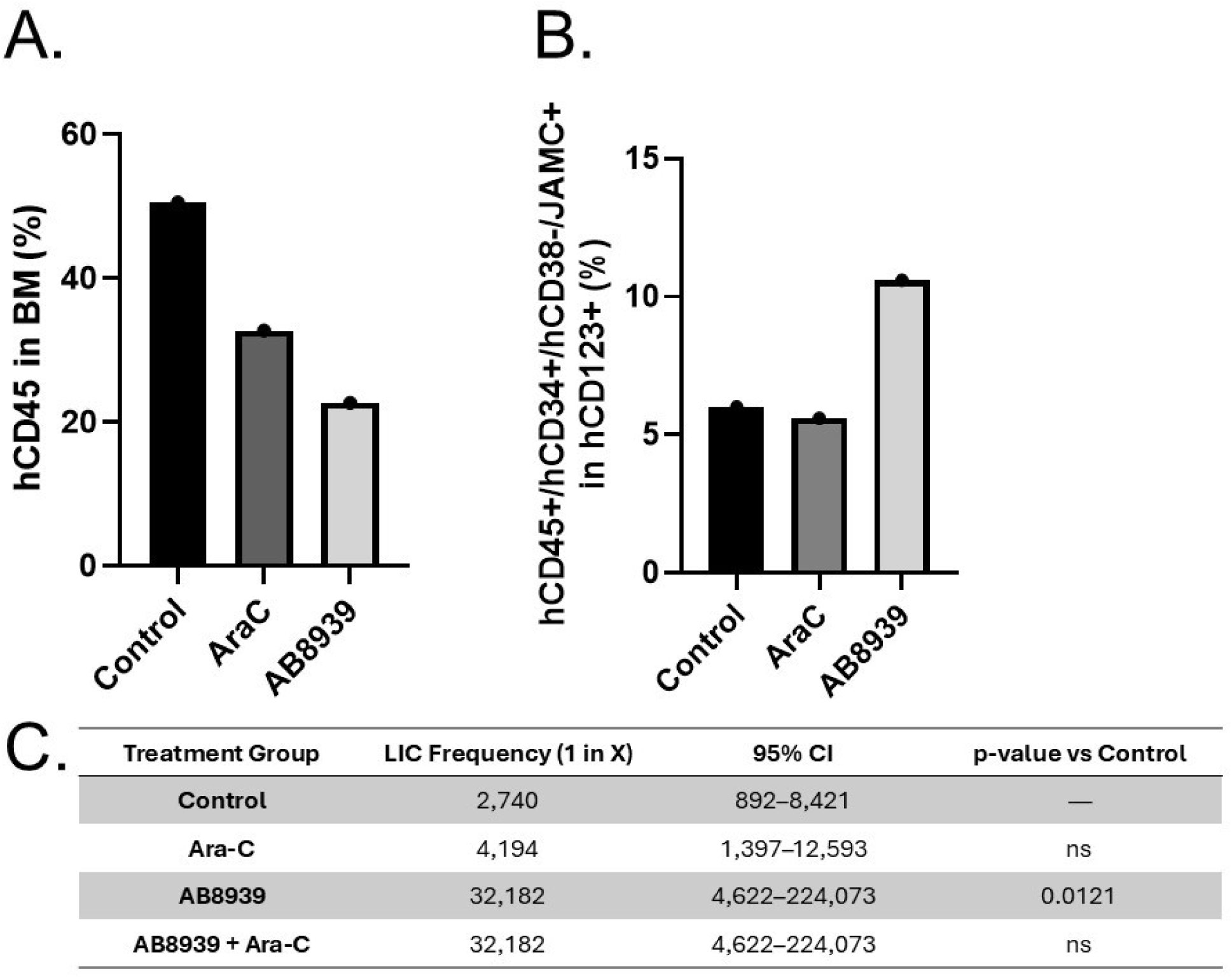
AB8939 targets Leukaemia inducing cells (LICs) and reduced leukaemia recurrence. NSG mice were engrafted with luciferase-tagged TG-LAM-36 AML PDX cells (AB8939-sensitive, Ara-C-resistant). After confirming engraftment, mice received 4 days of treatment: vehicle, Ara-C, AB8939, or AB8939 + Ara-C. Bone marrow was harvested 24 hours post-treatment for flow cytometric analysis of human leukemic cells (hCD45⁺) and LSC markers (CD34⁺/CD38⁻/CD123⁺/JAMC⁺). Panel A: AB8939 and Ara-C monotherapies reduced hCD45⁺ leukemic burden; AB8939 showed slightly greater efficacy. Combination therapy was comparable to AB8939 alone (***p < 0.001 vs. vehicle). Panel B: AB8939 increased the proportion of LSC-like cells within the hCD123⁺ population, likely due to selective blast depletion. Ara-C showed minimal LSC enrichment. Panel C: Extreme Limiting Dilution Analys (ELDA) revealed AB8939 (alone or in combination with Ara-C) reduced leukemia-initiating cell (LIC) frequency ∼12-fold vs. control (**p = 0.0121); Ara-C had no significant effect. Relative LIC frequency was calculated as Relative LIC frequency= LIC frequency_Control_/LIC frequency_Treatment_. These results highlight AB8939’s dual efficacy against bulk blasts and LSCs, supporting its potential to reduce AML relapse risk.

To assess the functional impact on leukemia-initiating cells (LICs), we performed Extreme Limiting Dilution Analysis (ELDA) on the re-transplanted mice. ELDA modeling revealed a dramatic reduction in LIC frequency following AB8939 treatment (1 in 32,182 cells; 95% CI: 4,622–224,073) compared to control (1 in 2,740 cells; 95% CI: 892–8,421; *p* = 0.0121). Ara-C alone had no significant effect (1 in 4,194 cells; 95% CI: 1,397–12,593), while the AB8939 + Ara-C combination mirrored the AB8939 monotherapy (1 in 32,182 cells; 95% CI: 4,622–224,073). Overall, ELDA confirmed significant differences in LIC frequency across groups (χ² = 10.4, df = 3, *p* = 0.0156), with AB8939 outperforming Ara-C and control in depleting functional LICs (Supplementary data, Tables S4 & S5). These findings demonstrate that AB8939 effectively targets the leukemia-initiating compartment in vivo, suggesting its potential to reduce relapse risk by eradicating residual LSCs that escape conventional therapy.

### Reverse Proteomics Revealed Aldehyde Dehydrogenases ALDH1s and ALDH2 as Main AB8939 Secondary Targets

To identify additional molecular targets that might account for the activity of AB8939 against leukemia-initiating cells (LICs), we employed a reverse proteomic strategy following the methodology described by Rix and colleagues (21). This approach involved generating an affinity matrix through covalent attachment of the compound to a resin. To enable this, we synthesized a series of AB8939 derivatives containing a primary amine (−NH₂) group, which allowed covalent cross-linking to NHS-activated resin for subsequent pull-down experiments using whole-cell extracts. Among these derivatives, AB9896—featuring an amino group appended to the terminal O-ethyl aliphatic chain (Figure 13)—emerged as the optimal candidate, as it preserved the full primary pharmacodynamic properties of AB8939.

**Figure 13.**
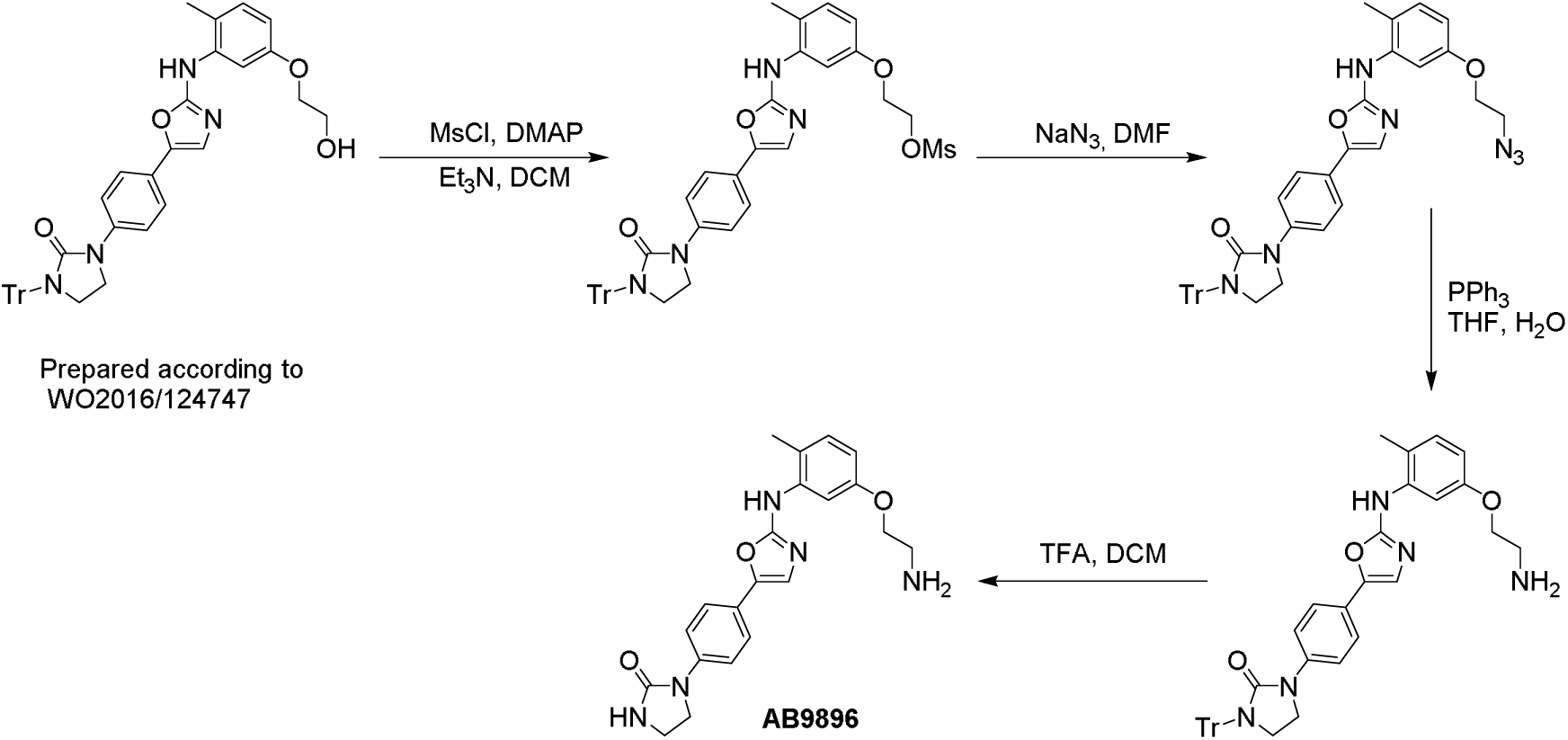
Synthesis of AB9896, a - NH2 derivative of AB8939. AB9896 is a functionalized derivative of AB8939 designed for reverse proteomics. A primary amine was introduced via a three-step synthesis to enable covalent coupling to NHS-activated resin for affinity chromatography. The modification preserved AB8939’s pharmacophore and biological activity, including antiproliferative potency, microtubule disruption, G2/M arrest, and tubulin polymerization inhibition. AB9896 was coupled to Sepharose beads for target capture from cell lysates; control beads were prepared using ethanolamine. Full synthesis and coupling protocols are detailed in Materials and Methods.

To maximize the likelihood of identifying novel AB8939 interactors, we first employed a mixed lysate prepared from AB8939-sensitive cell lines of diverse origins (hematopoietic, head and neck, ovarian, melanoma, etc.), thereby increasing protein diversity (Experiment 1). In a complementary approach, we used lysates derived from a single cell line, CLS354-4 (Experiment 2). Beads cross-linked with AB9896 were incubated with these lysates, and bound proteins were subsequently analyzed by LC–MS/MS and identified through protein database comparison. Relative protein abundance versus control and statistical significance were then calculated. The interactors identified through this reverse proteomic strategy are visualized in the volcano plot shown in Figure 14, which highlights significantly enriched proteins (red dots). Notably, members of the aldehyde dehydrogenase (ALDH) family were among the most enriched interactors, indicating a strong association between AB9896 and proteins involved in aldehyde metabolism and suggesting a potential functional link to this enzymatic pathway.

**Figure 14.**
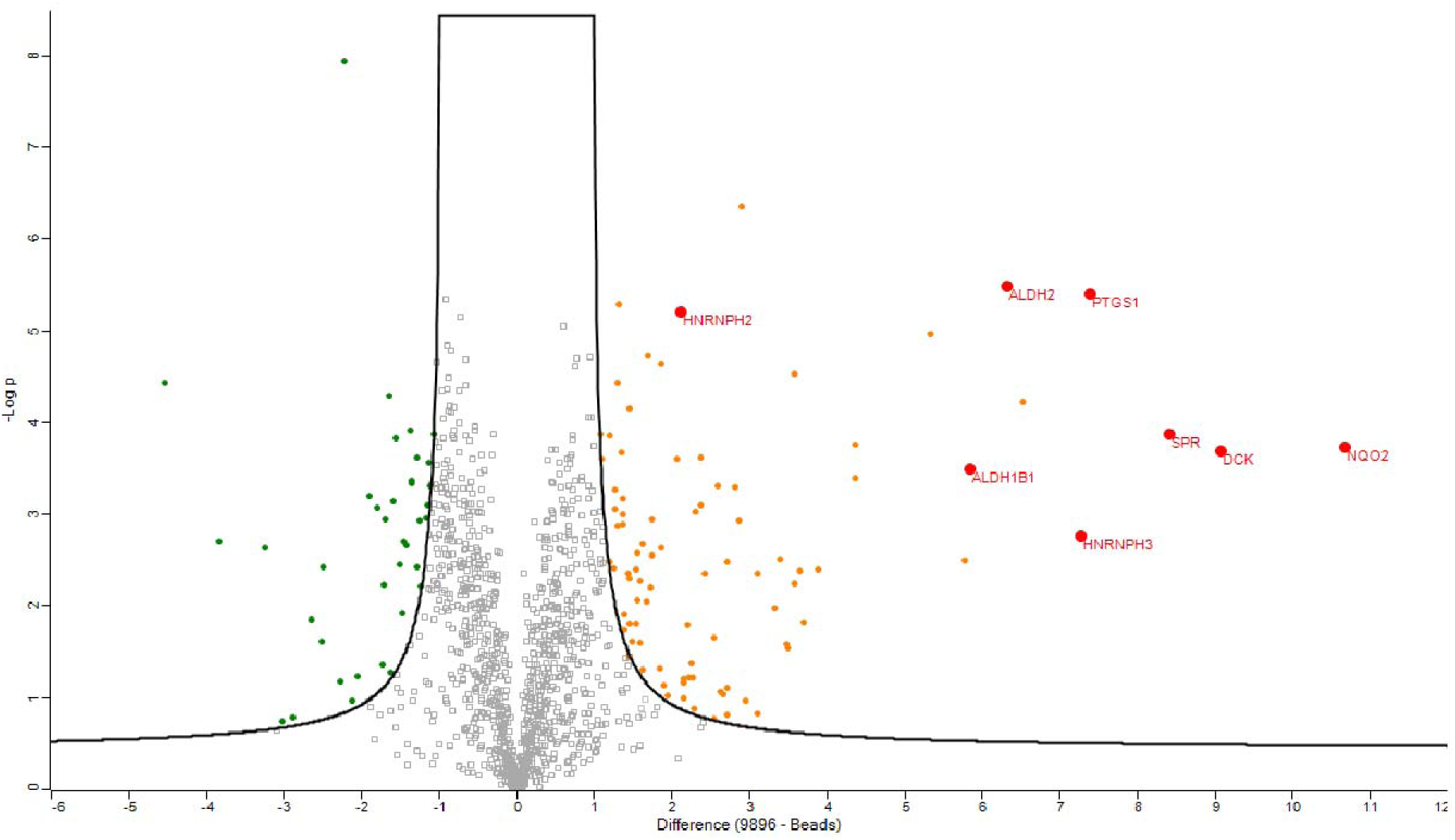
Volcano plot of Experiment 1 revealing ALDHs as main AB9896’s interactors. Volcano plot of affinity pull-down mass spectrometry (Experiment 1) to identify AB8939 protein targets. Lysates from 33 AB8939-sensitive tumor cell lines were incubated with AB9896-conjugated or control beads, and captured proteins were analyzed by LC-MS/MS with label-free quantification. The plot shows statistical significance (y-axis: –log₁₀ p-value) versus fold enrichment (x-axis: log₂ ratio of AB9896 vs. control). Proteins significantly enriched (fold change >2, p < 0.05) are highlighted in red. Major interactors include ALDH2 (34.5-fold, p < 0.0001), ALDH1B1 (7-fold, p < 0.01), and NQO2 (18-fold, p < 0.001). DCK (8-fold, p < 0.01) was enriched but not validated in enzymatic assays. Additional moderately enriched proteins (3–5 fold) included metabolic enzymes, chaperones, and cytoskeletal proteins.

The main ALDH isoforms interacting with AB9896, which were detected in both Experiment 1 (pool of 33 cell lysates) and Experiment 2 (CLS354-4 cell lysate), are summarized in Table 7 below.

**Table 7.** AB9896 pull down experiment identifies potential interactors.

| Accession | Gene | Peptide count |  |  |
| --- | --- | --- | --- | --- |
|  |  | AB9896 beads | Control | Ratio |

|  |  |  | beads | 9896/control |
| --- | --- | --- | --- | --- |
| **Experiment 1 (Pool of 33 cell lines lysates)** |  |  |  |  |
| P05091 | ALDH2 | 34,5 | 1 | 34,5 |
| P30837 | ALDH1B1 | 7 | 1 | 7 |
| **Experiment 2 (CLS354-4 cell lysate)** |  |  |  |  |
| P30837 | ALDH1B1 | 60 | 1 | 60 |
| P05091 | ALDH2 | 178 | 4 | 45 |
| P47895 | ALDH1A3 | 102 | 33 | 3 |

The most frequent interactor detected was NQO2 (Figure 14), a well-known promiscuous binder of kinase inhibitors, previously identified as a target of imatinib and dasatinib and potently inhibited by both drugs (21,22). The second most frequent interactor was deoxycytidine kinase (dCK); however, this is likely an artifact, as AB8939 showed no inhibitory activity against dCK in an in vitro kinase assay.

Of greater interest were members of the aldehyde dehydrogenase (ALDH) family—specifically ALDH1A3, ALDH1B1, and ALDH2 (Table 7). ALDH enzymes are increasingly recognized for their roles in tumor progression and therapy resistance. Elevated ALDH expression has been linked to enhanced proliferation, motility, and invasiveness across multiple cancer types, including breast, lung, colorectal, and prostate cancers (3). High ALDH activity supports malignant cell survival by promoting detoxification, thereby enabling resistance to apoptosis and facilitating metastasis. ALDH1A1-positive cancer stem-like cells, for example, display heightened tumorigenic potential and resistance to conventional therapies. Similarly, elevated ALDH2 expression has been associated with increased cancer cell survival, proliferation, and therapeutic resistance. Mechanistically, ALDH2 contributes to tumor maintenance by sustaining stem-like phenotypes, promoting epithelial–mesenchymal transition (EMT), and regulating oncogenic signaling pathways (4). Importantly, ALDH2 expression has also been identified as an adverse prognostic marker in AML post-remission outcomes (23).

### AB8939 Is a Potent Inhibitor of ALDH1A1 and ALDH2

We established an in vitro ALDH enzymatic assay using ALDH1A1 purified *in house* and commercially available ALDH2 (Abcam) enzymes, with activity monitored by NADH fluorescence at 340 nm. AB8939 inhibited both ALDH1A1 and ALDH2 with sub-micromolar potency (Figure 15), supporting its potential to target tumor cells with elevated ALDH activity, including cancer stem cell populations.

**Figure 15.**
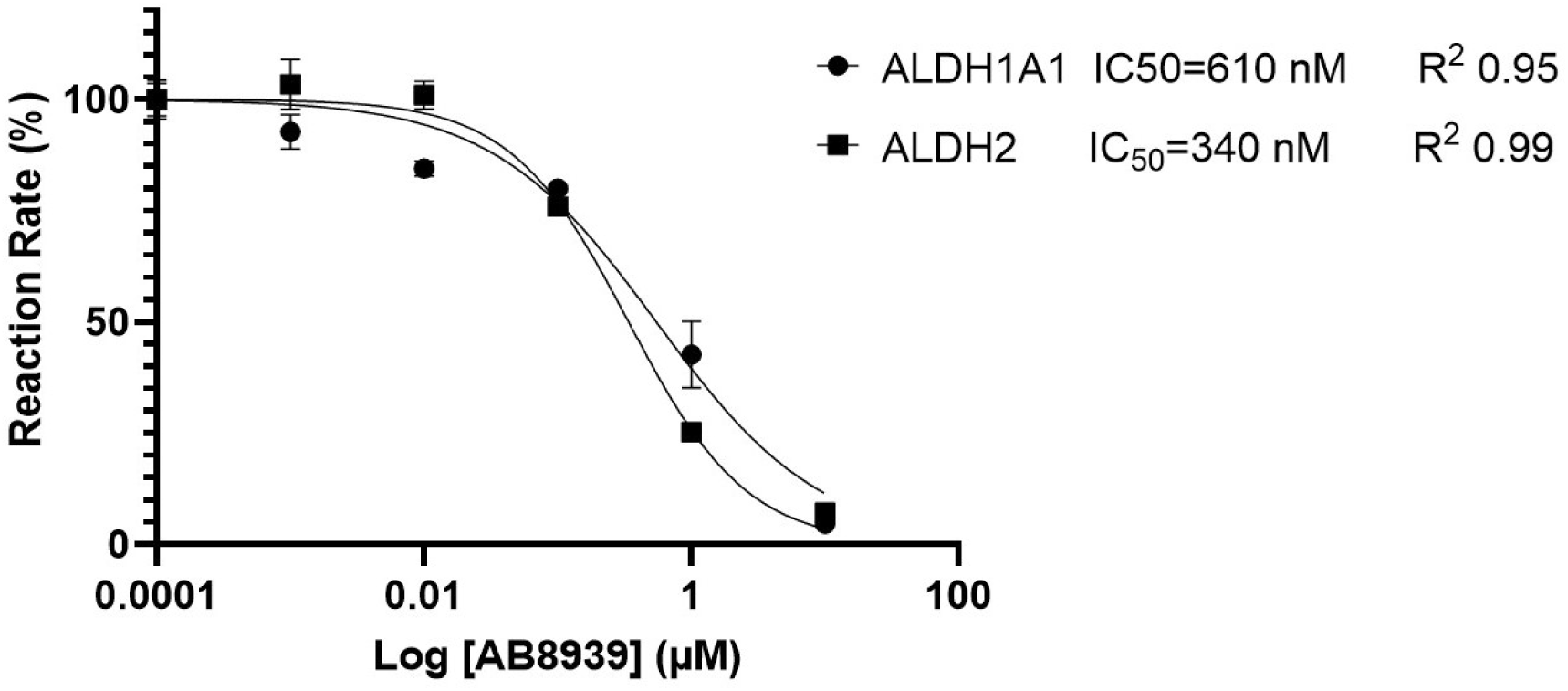
AB8939 inhibits recombinant ALDH1A1 and ALDH2 in vitro. Cell-free enzymatic assays using recombinant human ALDH isoforms with propionaldehyde substrate. Activity measured by NADH fluorescence (λ_ex=340 nm, λ_em=460 nm). IC₅₀ determined by nonlinear regression. Mean ± SEM from three technical triplicates.

In silico modeling indicated that, unlike the previously reported clinical ALDH1A1 inhibitor DIMATE, AB8939 binds with high affinity (reflected by low Gibbs free energy/binding energy) to the substrate-binding site of ALDH1A1 and ALDH2, rather than to the NAD⁺ site, as illustrated in Figure 16.

**Figure 16.**
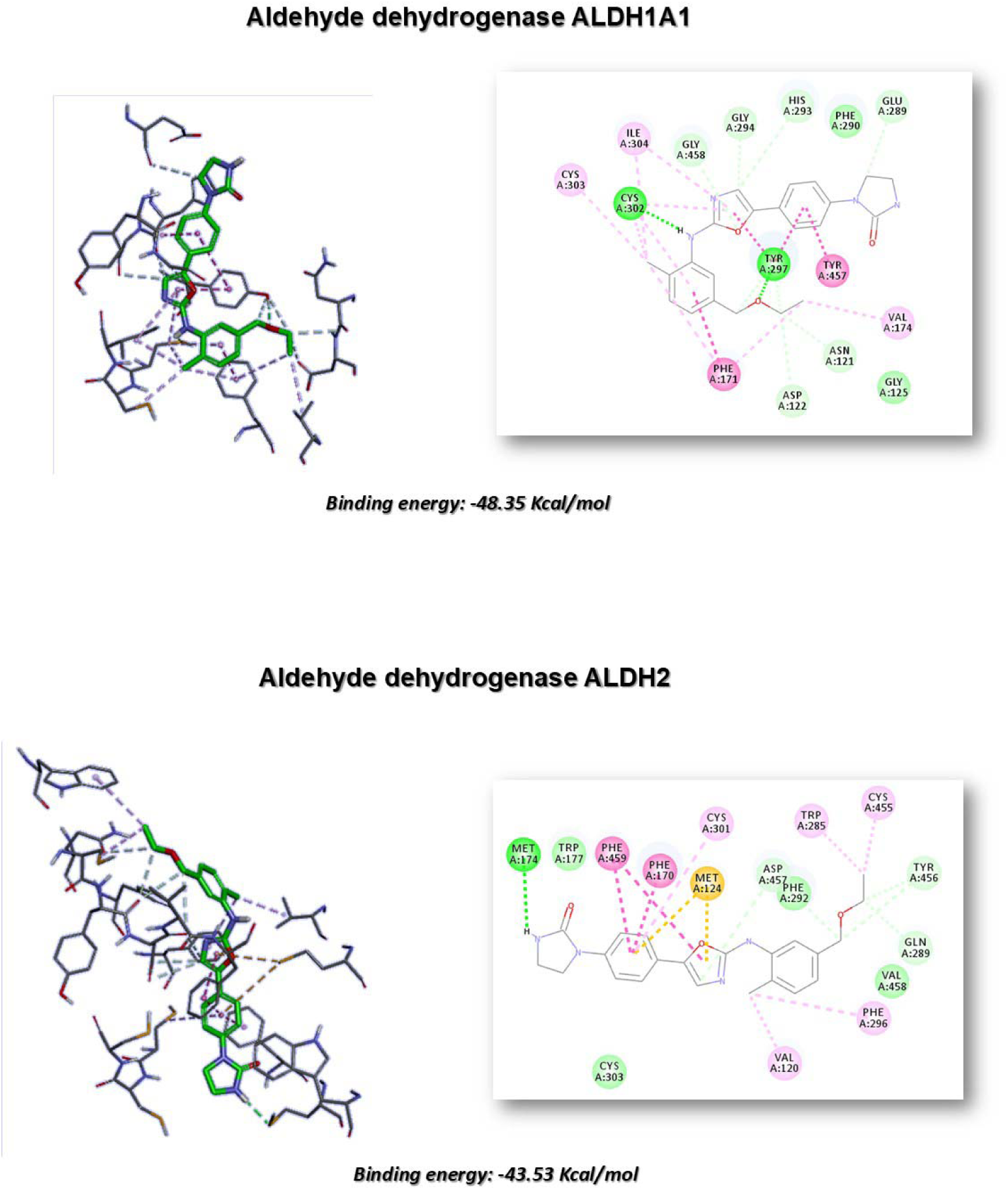
Proposed binding mode of AB8939 to the substrate site of ALDH1A1 and ALDH2. Molecular docking using Discovery Studio 2018 software package (DS; https://www.3ds.com/). Protein structures: ALDH1A1 (PDB 4WPN), ALDH2 (PDB 1O02). Panel A: AB8939 docked into ALDH1A1 substrate-binding tunnel. Docking score = −48.35 kcal/mol. AB8939 occupies substrate binding pocket, forming hydrogen bonds with Tyr297 and Cys302. Hydrophobic contacts with Phe171, Tyr457 (π-stacking), Ile304 and Cys303 (π-alkyl). Panel B: AB8939 in ALDH2 active site. Docking score = −43.53 kcal/mol. Hydrogen bonds with Met174, 169, Glu268, Thr244. π - stacking with Phe170 and Phe 459, π-sulfur with Met124 and π-sigma with Cys177, Trp285, Cys455, Val120 and Phe296. Overlaps with substrate binding site. Docking models predict AB8939 inhibits ALDH by binding the substrate/cofactor site. Predictions align with biochemical IC₅₀ values.

### AB8939 Inhibits ALDHs in Lung Tumor A549 Cells

To complement the findings obtained with recombinant ALDH enzymes, we next evaluated the functional activity of AB8939 using the Aldefluor cell-based assay. For this purpose, we selected the A549 lung adenocarcinoma cell line, which expresses high levels of all ALDH isoforms, as confirmed by Western blot analysis (Figure 17A). In addition to robust ALDH expression, A549 cells also display key markers of a cancer stem cell phenotype, including CD90 and CD133 (24). This makes A549 a relevant model for assessing the impact of AB8939 on tumor stem cell populations.

**Figure 17.**
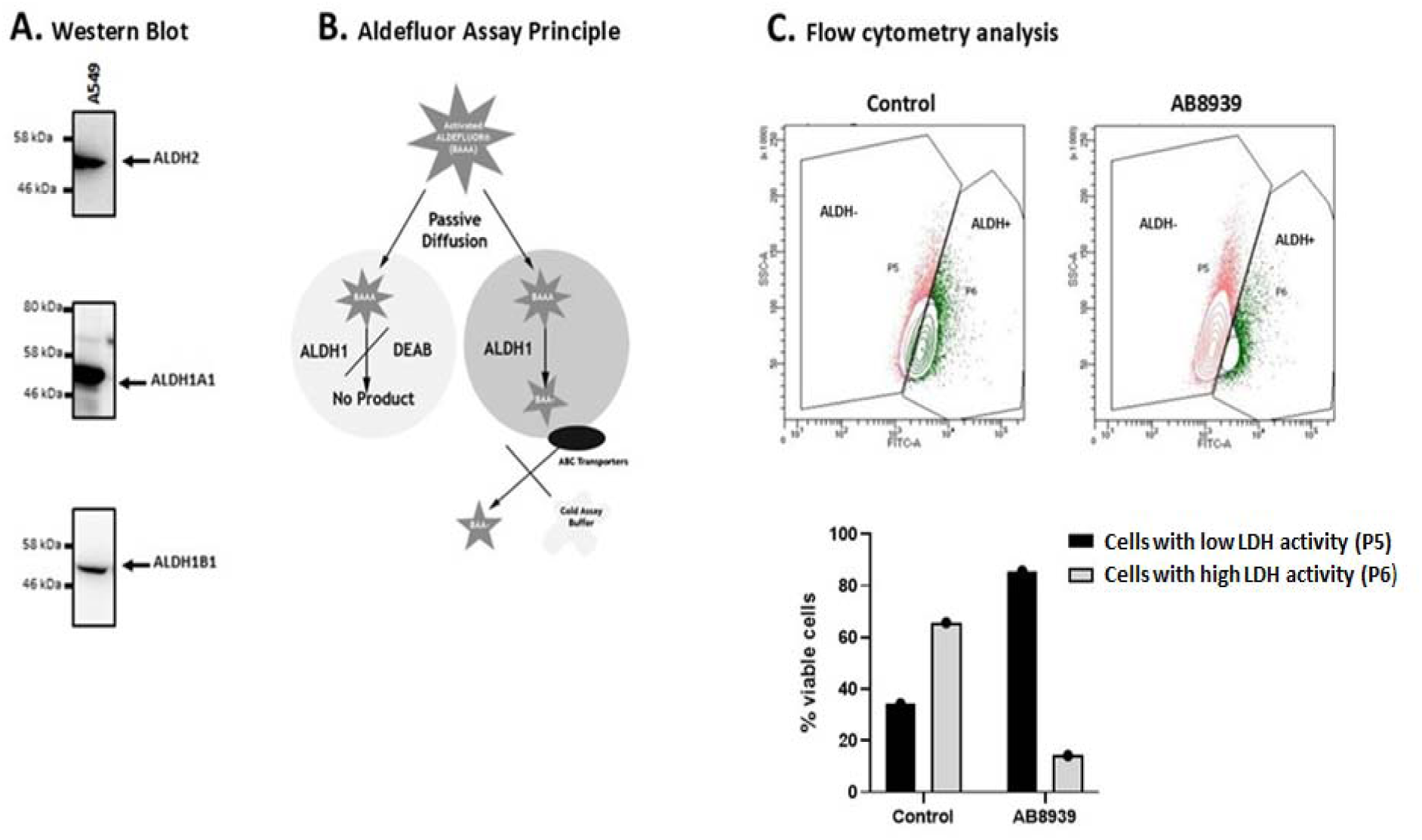
AB8939 inhibits ALDH activity in cells. Panel A: Western blot of ALDH isoforms in A549 lung adenocarcinoma cells. Cells express ALDH1A1 (∼55 kDa), ALDH1B1 (∼54 kDa), and ALDH2 (∼56 kDa). Panel B: Aldefluor assay schematic. BODIPY-aminoacetaldehyde (BAAA) enters cells and is converted by ALDH to fluorescent BODIPY-aminoacetate (BAA⁻), which accumulates intracellularly. ALDH inhibitors prevent conversion, reducing fluorescence. Panel C: Flow cytometry of A549 cells treated with vehicle, AB8939 (1 µM) for 3 hours, then incubated with Aldefluor substrate for 40 minutes. Vehicle: 65% ALDH^bright^ cells. AB8939: 10% ALDH^bright^ (85% reduction, p<0.001 vs vehicle). AB8939 effectively inhibits ALDH activity in live cancer cells, supporting potential to target ALDH^high^ cancer stem cell populations. Dual targeting of microtubules and ALDH positions AB8939 as a multi-targeted therapeutic agent.

The Aldefluor reagent system enables detection of intracellular ALDH activity through a fluorescent substrate. It is supplied as Bodipy-aminoacetaldehyde diethyl acetal (BAAA-DA), which is converted under acidic conditions to Bodipy-aminoacetaldehyde (BAAA), a fluorescent substrate for ALDH (Figure 17B). In this study, A549 cells were treated with either DMSO (control) or AB8939 (1 µM, 3 hours), followed by Aldefluor labeling in the presence of the inhibitor for 40 minutes and subsequent flow cytometry analysis (Figure 17C, upper panel). AB8939 treatment markedly reduced the proportion of Aldefluor-positive cells, as shown in Figure 17C (lower panel), demonstrating potent inhibition of cellular ALDH activity. Consistently, flow cytometry histograms revealed a sharp decline in the fraction of A549 cells with active ALDH, from 65% in controls to 10% following AB8939 exposure (Figure 17C). These results confirm that AB8939 effectively inhibits ALDH catalytic activity in living cancer cells.

## Discussion

We report the identification of AB8939 as a novel small-molecule microtubule-disrupting agent (MDA) with potent and broad-spectrum antiproliferative activity. Across diverse tumor cell lines, hematopoietic malignancies were particularly sensitive, with most lines inhibited after 72 hours of treatment. Prolonged exposure further demonstrated that AB8939 effectively eliminated all tested tumor types, including those harboring oncogenic driver mutations or displaying cancer stem cell–like phenotypes, such as A549 lung cancer cells.

In biochemical assays, AB8939 strongly inhibited tubulin polymerization and disrupted preformed microtubules. Treated cells exhibited disorganized microtubule networks and punctate cytoplasmic staining consistent with depolymerization and accumulation of free tubulin. X-ray crystallography confirmed binding of AB8939 to the colchicine-binding site of tubulin. Similar to other colchicine-site ligands (e.g., VERU-111), AB8939 overlaps closely with colchicine but does not engage the deeper β-subunit subsite (25). Like many compounds in this class (26), AB8939 primarily establishes hydrophobic interactions, with limited hydrogen bonding. Notably, polar contacts with Asp251 and Asn349 mirror those stabilizing VERU-111, either directly (Asp251) or via a water molecule (Asn349). Collectively, these findings establish AB8939 as a bona fide colchicine-site binder.

Targeting the tubulin network remains a validated therapeutic strategy, as exemplified by the vinca alkaloids vinblastine and vincristine, both FDA-approved for multiple malignancies. However, colchicine itself is unsuitable as an anticancer agent due to dose-limiting toxicities, including gastrointestinal injury, multi-organ failure, and severe adverse events such as neuropathy, myelosuppression, and cardiotoxicity (27). Moreover, the clinical efficacy of microtubule-targeting agents (MTAs) is frequently compromised by resistance mechanisms, most notably drug efflux mediated by P-glycoprotein (P-gp, MDR1). P-gp reduces intracellular drug accumulation and confers resistance not only to MTAs (e.g., paclitaxel, docetaxel, vincristine) but also to anthracyclines (e.g., daunorubicin, doxorubicin, mitoxantrone, idarubicin).

Strikingly, AB8939 is not a P-gp substrate and retains high potency in P-gp–overexpressing cells resistant to paclitaxel, vincristine, or doxorubicin. This property is particularly relevant for older AML patients (≥55–60 years), who often cannot tolerate intensive chemotherapy and whose leukemias frequently harbor poor-prognosis karyotypes and express P-gp (28). Expression of multidrug resistance markers has also been linked to immature, stem-like leukemia phenotypes that contribute to relapse. Current therapeutic strategies in this population remain largely ineffective, with overall survival rates below 10%.

Beyond P-gp–mediated resistance, overexpression of class III β-tubulin has been implicated in resistance to MTAs, including taxanes (29). AB8939 demonstrated robust activity in β3-tubulin–high cell lines such as A549, H1299, PANC-1, and MES-SA. Importantly, AB8939 also displayed strong ex vivo cytotoxicity against blasts from AML patients, including refractory and relapsed cases. It was active against blasts resistant to vincristine and cytarabine (Ara-C), and showed marked efficacy in poorly differentiated AML-M1, AML secondary to MDS, and monocytic AML subtypes. Notably, AB8939 was effective not only against treatment-naïve AML blasts but also against relapsed and refractory samples, underscoring its potential as a therapeutic option for high-risk AML.

In summary, AB8939 emerges as a dual-acting agent, combining potent tubulin destabilization with inhibition of aldehyde dehydrogenases (ALDHs). This unique pharmacological profile may provide a two-pronged therapeutic benefit: direct cytotoxicity through disruption of microtubule dynamics, and functional impairment of ALDH-driven resistance pathways that sustain leukemia stem cells and contribute to relapse. By simultaneously targeting proliferative blasts and stem-like subpopulations, AB8939 holds promise not only for overcoming chemoresistance but also for achieving deeper and more durable remissions in AML. Reflecting this potential, AB8939 is now being evaluated in a Phase I clinical trial in combination with venetoclax in patients with relapsed or refractory AML (NCT05855300). Future studies dissecting the relative contribution of ALDH inhibition to AB8939’s efficacy will be critical to fully exploit this dual mechanism in clinical development.

## MATERIALS AND METHODS

### Tumor Cells lines and culture

All cell culture media were obtained from Life Technologies and supplemented with either L-glutamine or GlutaMAX, 100 U/mL penicillin and 100 µg/mL streptomycin (GIBCO, ref. 15140122), and 10% (v/v) heat-inactivated fetal calf serum (FCS; AbCys, lot S02823S1800). For MEM medium, additional supplements included 1% non-essential amino acids (GIBCO, ref. 11140035) and 1 mM sodium pyruvate (GIBCO, ref. 111360070), in addition to GlutaMAX and 10% FCS. All cell lines were maintained under standard culture conditions (37 °C, humidified atmosphere of 5% CO₂ in air).

| Tumor type | Cell line | Culture medium* | Gift/Purchase origine |
| --- | --- | --- | --- |
| Breast Cancer | BT-20, BT-474, MCF-7, HCC1-937, MDA-MB-134, BRCAMZ-01 | RPMI | Dr Dubreuil, CRCM U1068 INSERM, Marseille |
|  | MDA-MB-231 | DMEM : F12 | Dr Dubreuil, CRCM U1068 INSERM, Marseille |
| Colorectal Cancer | DLD-1, HRT-18 | RPMI |  |
|  | HCT116, CACO-2 | MEM |  |
|  | HT-29, SW-480, | DMEM |  |
|  | LS174T |  |  |
| **Gastric Cancer** | AGS, HGC-27 | DMEM, DMEM : F12 | Cell Line Service, Germany |
| **Glioblastoma** | U87MG, U118 | DMEM : F12, MEM | Cell Line Service, Germany |
| **Head and Neck Cancer** | A64CLS, CLS-354-4, COLO680N | RPMI | Cell Line Service, Germany |
|  | Detroit 562 | DMEM : F12 | Cell Line Service, Germany |
|  | Hep2, RPMI2650, SW579 | MEM | Cell Line Service, Germany |
| **Hepatoma** | Hep G2 | DMEM | Cell Line Service, Germany |
|  | PLC PRF5 | DMEM : F12 | Cell Line Service, Germany |
| **Melanoma** | A375, LU1205, M6, M67, MelSol, MelWO, MelSek28 |  | Dr. Tartare-Deckert INSERM U895, Nice |
| **NSCLC** | A549, Calu-6, GCT, H1299, H1650, H820, HCC827 |  | Dr Dubreuil, CRCM U1068 INSERM, Marseille |
| **Ovarian Cancer** | MES-SA, OV90, OVCAR-3, TOV112D, TOV21G, O114 |  | Dr Dubreuil, CRCM U1068 INSERM, Marseille |
| **Prostate Cancer** | DU145, LNCAP, PC-3 |  | Dr. Juan Iovanna, CRCM U1068 INSERM, Marseille |
| **Pancreatic Cancer** | BxPC-3, Capan-2, PANC-1, MiaPaca |  | Dr. Juan Iovanna, CRCM U1068 INSERM, Marseille |
| **Lymphoma T Cell** | CCRF-CEM, HUT-78, JURKAT, MOLT-13, PEER, RPMI 8401 | RPMI | Dr. Olivier Hermine, Institut Imagine INSERM U1163, Paris |
| **Lymphoma B Cell** | HLB-2, JEKO-1, JVM-3, MEC-1, RAJI, REC-1, RL, SUDHL-4, RAMOS | RPMI | Dr. Olivier Hermine, Institut Imagine INSERM U1163, Paris |
| **AML** | HL-60, MOLM-14, MOLM-13, NB4, NOMO-1, THP-1, RS4-11, U-937, MV-4-11 | RPMI | Dr Dubreuil, CRCM U1068 INSERM, Marseille |
| **AMLK** | AML ETO2-GLIS2, CHRF, CMK | RPMI | Dr. Thomas Mercher (Institut Goustave Roussy, Paris) |

### Cell Cycle Analysis

#### Method 1

Cells were seeded in 24-well plates at a density of 1 × 10LJ cells/mL per well in their respective culture medium. After 24 hours, the medium was replaced with fresh medium containing AB8939 at 10, 100, or 1000 nM, as indicated. Following 24 hours of treatment, cells were washed once with 1× PBS and detached using 0.05% trypsin–EDTA (Life Technologies, cat. #25300-054). Cells were resuspended in medium, transferred to 96-well plates, centrifuged, and washed twice with cold PBS. Pellets were then resuspended in propidium iodide (PI) staining solution containing 0.1% NP-40, 0.1% sodium citrate, 50 µg/mL PI, and 0.2 mg/mL RNase A. DNA content was analyzed immediately using an Accuri C6 flow cytometer.

#### Method 2

Cell cycle distribution was also assessed using the Muse™ Cell Cycle Kit (Millipore, cat. #MCH100106) according to the manufacturer’s instructions. Briefly, cells were detached with 0.05% trypsin–EDTA, resuspended in medium, transferred to 1.7 mL tubes, centrifuged at 300 g for 5 minutes, and washed once with 1× PBS. Pellets were fixed in ice-cold 70% ethanol and incubated for at least 3 hours at −20 °C. After centrifugation (300 g, 5 minutes) and one PBS wash, 200 µL of Muse Cell Cycle reagent was added to each pellet. Samples were incubated for 30 minutes at room temperature in the dark and analyzed on the Muse® Cell Analyzer (Merck Millipore).

### Annexin V Apoptosis Assay

Apoptosis was quantified using the Muse™ Annexin V & Dead Cell Assay (Millipore, cat. #MCH100105) following the manufacturer’s protocol. This assay employs Annexin V to detect externalized phosphatidylserine on apoptotic cells, together with a dead cell marker that discriminates between intact and permeabilized membranes. Briefly, cells were detached with 0.05% trypsin–EDTA, collected in medium, transferred to 1.7 mL tubes, centrifuged at 300 g for 5 minutes, and resuspended in 100 µL of culture medium. An equal volume (100 µL) of Muse Annexin V & Dead Cell reagent was added, and samples were incubated for 20 minutes at room temperature in the dark before analysis on the Muse® Cell Analyzer (Merck Millipore).

### Cell-Based Proliferation and Viability Assay

Cell proliferation and viability were assessed using the CellTiter-Blue® assay (Promega, cat. #G8081). Briefly, 1 × 10LJ cells per well were seeded in 50 µL of medium in 96-well plates. After 24 hours, treatment was initiated by adding 2× drug solutions prepared as serial dilutions (½ or ⅓) over a concentration range of 0–10 µM. Following 72 hours of incubation at 37 °C, 10 µL of CellTiter-Blue reagent was added to each well, and plates were incubated for an additional 4 hours.

Fluorescence intensity, proportional to the number of viable cells, was measured at 544 nm excitation and 590 nm emission using a POLARstar OMEGA microplate reader (BMG Labtech). Wells without cells served as background controls, while untreated cells were used as positive controls (defined as 100% proliferation). Each condition was tested in duplicate, and experiments were independently repeated at least twice.

Data were exported to Excel, where mean values and standard deviations were calculated and expressed as a percentage of untreated controls. All proliferation datasets were archived in AB Science’s corporate database (Dotmatics platform).

### Ex Vivo AML Cell Viability Assay

Peripheral blood (PB) and/or bone marrow (BM) samples from AML patients were collected at diagnosis or relapse after informed consent (HematoBio, NCT05602168; CeGAL, NCT02619071; Department of Hematology, Institut Paoli-Calmettes, France). For viability assays, Ficoll-purified mononuclear cells from AML patients or patient-derived xenografts (PDXs; generated by pooling bone marrow and spleen cells from transplanted mice) were seeded in 96-well plates in complete medium (RPMI supplemented with 20% fetal calf serum). Cells were exposed for 48 hours to increasing concentrations of AB8939, cytarabine (Ara-C), or vincristine.

Cell viability was quantified using the CellTiter-Glo® luminescent assay (Promega, cat. #G7571). Briefly, 50 µL of reagent was added per well, followed by 2 minutes of orbital shaking and a 10-minute incubation at room temperature. Luminescence was measured with a Centro XS3 LB 960 Microplate Luminometer (Berthold Technologies).

Data were expressed as IC₅₀ values (µM). For comparative analyses of AB8939 versus vincristine, relative IC₅₀ values were calculated by normalizing to the predefined IC₁₀₀ of each drug (2 µM for vincristine; 5 µM for AB8939), corresponding to the concentration inducing complete cell death in the assay.

### Immunofluorescence Staining of Microtubule and Actin Networks

Eight-well Lab-Tek chamber slides (ref. 177402) were pre-coated with fibronectin (10 µg/mL). Approximately 5 × 10LJ cells were seeded per well and incubated overnight at 37 °C in a humidified atmosphere containing 5% CO₂. The following day, the medium was replaced with fresh medium containing 100 nM AB8939. After 1–3 hours of treatment, cells were washed twice with PBS and fixed for 10 minutes at room temperature in 4% formaldehyde. Following two additional PBS washes, cells were permeabilized in PBS supplemented with 3% BSA and 0.1% Triton X-100.

Actin filaments were stained with Phalloidin-FITC (Life Technologies, cat. #A12379) diluted 1:40 in PBS/3% BSA/0.1% Triton X-100 for 45 minutes at room temperature. Microtubules were stained for 1 hour at room temperature with a rat monoclonal anti–α-tubulin antibody (Life Technologies, cat. #MA1-80189) diluted 1:500 in PBS/3% BSA/0.1% Triton X-100, followed by two PBS washes and incubation with Alexa Fluor 594–conjugated anti-rat secondary antibody (Abcam, cat. #150160) diluted 1:1000 for 45 minutes at room temperature.

After staining, cells were washed 3–4 times with PBS and counterstained with DAPI (Sigma, cat. #D8417; 1:5000 dilution). Slides were analyzed using the EVOS Cell Imaging System (Thermo Fisher Scientific).

### In Vitro Tubulin Polymerization Assay

Tubulin polymerization was assessed using the fluorescence-based Tubulin Polymerization Assay Kit (Cytoskeleton, cat. #BK011P) according to the manufacturer’s instructions. Briefly, purified porcine brain tubulin was prepared at 2 mg/mL in polymerization buffer (80 mM PIPES, pH 6.9; 2.0 mM MgCl₂; 0.5 mM EGTA; 1.0 mM GTP; 15% glycerol) and kept on ice. Tubulin was then incubated at 37 °C with varying concentrations of AB8939 or control compounds in the presence of 1 mM GTP.

Polymerization was monitored by fluorescence enhancement resulting from incorporation of a fluorescent reporter into microtubules during assembly. Experiments were performed in duplicate, with paclitaxel and nocodazole included as positive controls for polymerization and depolymerization, respectively.

### Crystallization, Data Collection, and Structure Refinement of AB8939–Tubulin Complex

Crystals of subtilisin-treated tubulin in complex with the stathmin-like domain of RB3 (5) were soaked in crystallization buffer supplemented with 250 µM AB8939 and subsequently flash-cooled in liquid nitrogen. X-ray diffraction data were collected at the PROXIMA 2A beamline (SOLEIL Synchrotron) and processed using autoPROC (6), which incorporates the STARANISO procedure for anisotropy correction (http://staraniso.globalphasing.org/).

The previously solved sT2R structure (PDB ID: 3RYC) served as the starting model. Refinement was performed with BUSTER (7) version 2.10.4 (Global Phasing Ltd) in combination with iterative model building in Coot (8). Data collection and refinement statistics are provided in Table S2.

The final atomic coordinates and structure factors have been deposited in the Protein Data Bank under accession code 8CGZ. Structural figures were generated using PyMOL (www.pymol.org).

### P-gp-Glo Assay

The effect of test compounds on P-glycoprotein (P-gp) activity was evaluated using the P-gp-Glo™ assay system (Promega). This assay distinguishes between P-gp substrates, which stimulate ATPase activity, and inhibitors, which reduce ATP consumption in the presence of a known substrate. None of the compounds tested acted as P-gp inhibitors.

Compounds were first dissolved in DMSO at 40× working concentration and subsequently diluted to 2.5× in P-gp assay buffer, ensuring a final DMSO concentration of 0.5% in the reaction mixture. Reactions were performed in 96-well half-area white plates (Corning Costar, ref. 3693). Each well contained 20 µL of compound solution mixed with 20 µL of 2.5× recombinant P-gp (2.5 mg/mL; 25 µg). After a 5-minute pre-incubation at 37 °C, reactions were initiated by adding 10 µL of 25 mM Mg²⁺-ATP (final concentration 5 mM) and incubated for 40 minutes at 37 °C.

Reactions were terminated by adding 50 µL of ATP detection reagent per well, followed by 20 minutes of incubation at room temperature. Luminescence was measured using a Clariostar plate reader (BMG Labtech). Raw data were processed according to the manufacturer’s instructions and analyzed with GraphPad Prism® v10.6.1., applying the Michaelis–Menten model.

### AB9896 Pull-Down Assay

AB8939-NH₂ (hereafter referred to as AB9896) was covalently coupled via its amino group to NHS-activated Sepharose 4 Fast Flow beads, following the manufacturer’s instructions (Cytiva Life Sciences). Pull-down experiments were performed using either a pooled lysate from 33 cell lines (first experiment) or a CLS354-4 cell lysate (second experiment).

Cells were cultured to ∼90% confluence and lysed in HNTG buffer (50 mM HEPES, pH 7.0; 150 mM NaCl; 1% Triton X-100; 10% glycerol) supplemented with 100 µM sodium orthovanadate and a protease inhibitor cocktail (Complete Mini, EDTA-free; Roche Diagnostics). Lysates (4 mL) were pre-cleared with uncoupled NHS-activated beads for 4 hours at 4 °C, then incubated overnight at 4 °C with either control beads or AB9896-coupled beads (1 mL lysate with 200 µL beads). Beads were washed three times with HNTG buffer, resuspended in sample buffer, and stored at −20 °C until further analysis.

For quality control, 10% of bead-bound proteins were separated on NuPAGE 4–12% Bis-Tris gels (Invitrogen) in MOPS buffer and visualized by silver staining. For mass spectrometry, the remaining 90% of pull-down proteins were resolved on NuPAGE gels, stained with Imperial Blue (Thermo Scientific), excised, reduced with DTT, alkylated with iodoacetamide, and digested with sequencing-grade trypsin (Promega). Extracted peptides were concentrated prior to LC–MS/MS analysis.

### Mass Spectrometry

Mass spectrometry analyses were performed by LC–MS/MS using an LTQ-Velos-Orbitrap mass spectrometer (Thermo Electron) coupled to a nanoLC Ultimate 3000 Rapid Separation system (Dionex). A 5 µL injection volume, corresponding to 20% of the total sample, was loaded onto a Dionex Acclaim PepMap 100 pre-column (C18, 2 cm × 100 µm i.d., 100 Å pore size, 5 µm particle size) for pre-concentration and washing. Peptides were then separated on a Dionex Acclaim PepMap RSLC analytical column (C18, 15 cm × 75 µm i.d., 100 Å pore size, 2 µm particle size) at a flow rate of 300 nL/min using a two-step linear gradient: 4–20% acetonitrile/0.1% formic acid (90 min), followed by 20–45% acetonitrile/0.1% formic acid (30 min). Separation was monitored by UV absorbance at 214 nm.

Peptides were ionized in a nanospray source with a spray voltage of 1.4 kV and a capillary temperature of 275 °C. Data were acquired in data-dependent acquisition mode. Survey full scans were recorded in the Orbitrap (m/z 300–1700) at 30,000 FWHM resolution (at m/z 400), with a target AGC value of 1 × 10LJ and a maximum injection time of 500 ms. In parallel, the 10 most intense precursor ions were selected for CID fragmentation and analyzed in the linear ion trap (normalized collision energy 35%, activation time 10 ms, AGC target 1 × 10LJ, maximum injection time 100 ms, isolation window 2 Da). Parent ion masses were automatically calibrated using the 445.120025 m/z lock mass.

Fragment ions were measured in the linear ion trap to maximize sensitivity and MS/MS coverage. Dynamic exclusion was applied with a repeat count of 1 and an exclusion duration of 30 s. Each sample was analyzed in triplicate, with blank runs performed prior to each analysis to monitor background.

### Proteomic Analysis

Raw mass spectrometry files were processed using Proteome Discoverer 1.3 (Thermo Fisher Scientific) and searched via an in-house Mascot server (version 2.4.1; Matrix Science Inc.) against the human subset of the SwissProt database (20,264 sequences; release 2014-03). Database searches were performed with the following parameters: maximum of one trypsin missed cleavage, methionine oxidation as a variable modification, cysteine carbamidomethylation as a fixed modification, peptide mass tolerance of 6 ppm, and fragment mass tolerance of 0.8 Da. Only peptides with high Mascot scores and a false discovery rate (FDR) <1% were retained for protein identification. Pull-down experiments were performed with both drug-coupled and control beads to discriminate specific interactors from nonspecific contaminants. A complete list of identified proteins is provided in the supplementary Excel file (*proteomic.xlsx*).

For relative quantification, label-free intensity-based analysis was performed using Progenesis QI for Proteomics v1.0 (Nonlinear Dynamics) according to the manufacturer’s instructions. Raw LC–Orbitrap MS data were imported to generate LC–MS heatmaps of retention time versus m/z. Features were aligned and filtered using the following criteria: retention time 15–135 min, m/z 300–1700, charge state 2–6, and ≥3 isotopes. MS/MS spectra were exported as Mascot Generic Files (MGF), selecting the three most intense precursors per feature with precursor intensity >25%.

MGF files were searched against the human SwissProt subset (release 2014-03) using Mascot with the following parameters: one missed cleavage allowed, methionine oxidation and cysteine carbamidomethylation as fixed modifications, peptide mass tolerance of 6 ppm, and fragment mass tolerance of 0.8 Da. Peptides were filtered at 5% FDR with an ion score cutoff of 20 before re-import into Progenesis for protein grouping and quantification.

Protein quantification was based on normalized total ion intensities of all non-conflicting peptides. Conflicting peptide identifications were excluded. Statistical analysis was performed within Progenesis using one-way ANOVA, with p-values calculated from the sum of normalized peptide abundances across all runs.

### Molecular Docking of AB8939 in ALDH1A1 and ALDH2

#### Protein preparation

The crystal structure of the binding complex, was downloaded from the RCSB Protein Data Bank. Water molecules and co-crystallized ligands were deleted and then the receptor protein was subsequently prepared by the Prepare Protein protocol using the Discovery Studio 2018 software package (DS; https://www.3ds.com/). The following tasks were performed: insertion of missing atoms in incomplete residues, modeling of missing loop regions, deleting of alternate conformations (disorder), standardizing of atom names, and protonating of titratable residues.

#### Molecular docking

The protein was prepared by removing the water, adding hydrogen, insertion of missing atoms in incomplete residues, modeling of missing loop regions, deleting of alternate conformations (disorder), standardizing of atom names, and protonating of titratable residues correcting. Then, CFF charges and potentials were assigned to the protein with CHARMm27 force field. The active site was predicted and identified using “Edit binding site” module of DS2018 according to the native ligand and the radius was set to 10Å. The final poses were saved for each ligand based on scoring and ranking by the negative value of GOLD score energy. The remaining parameters were default

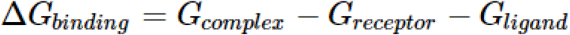

The binding free energy (Δ*G*_bind_) was calculated with the molecular mechanics and Poisson–Boltzmann solvation area (MM/PBSA) methodology was applied based on stable interactions. Entropy was omitted herein because we pay more attention to relative binding free energy for a series of very similar systems.

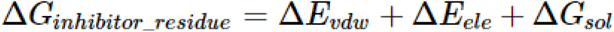

Where *G_complex_*, *G_receptor_*, and *G_ligand_*are the free energy of complex, receptor and ligand molecules, respectively. Then, free energy of binding for a receptor-ligand complex can be calculated from the free energies of the complex, the receptor, and the ligand. Using CHARMm based energies and implicit solvation methods it is possible to estimate these free energies and thus calculate an estimate for the overall binding free energy. MM-PBSA was used to compute the interaction energies to each pose complex in CHARMM27 by considering molecular mechanics energies and solvation energies without considering the contribution of entropies. The binding interaction between residue-ligand pair comprises three terms: the van der Waals contribution (Δ*E_vdw_*), the electrostatic contribution (Δ*E_ele_*) and the solvation contribution (Δ*G_sol_*).

#### In Vitro ALDH Activity

Human full-length recombinant ALDH1A1 was expressed and purified in-house in *E. coli* as previously described (X. Wang & Weiner, 1995). Enzyme purity (>95%) was confirmed by SDS–PAGE with Coomassie Blue staining. Fractions containing ALDH activity were pooled and concentrated using a Centricon centrifugal filter device (Amicon). Purified enzyme was stored at −80°C in 50% glycerol until use. Recombinant human ALDH2 (AA 17–517) was purchased from Abcam (cat. #ab87415).

ALDH dehydrogenase activity assays were performed in 384-well plates (Greiner Bio-One, ref. 781080) using 25 mM sodium HEPES buffer (pH 7.5) at room temperature. Enzymatic activity was monitored by measuring NADH fluorescence over time with a Pherastar FS plate reader (BMG Labtech). Reactions were initiated by adding 20 µL of a 2× acetaldehyde substrate/AB8939 mixture to 20 µL of 2× enzyme buffer containing ALDH (60 nM) and NAD⁺ (500 µM for ALDH1A1; 400 µM for ALDH2). Fluorescence emission at 460 nm was recorded, and reaction velocities were calculated for each AB8939 concentration. Inhibition curves were generated in GraphPad Prism using the variable slope equation:

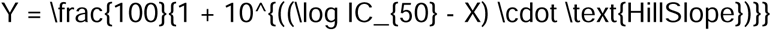

#### Aldefluor Assay

Cellular ALDH activity was assessed in A549 lung adenocarcinoma cells using the Aldefluor reagent system (Stemcell Technologies, cat. #01703), according to the manufacturer’s instructions. The assay is based on the conversion of BODIPY-aminoacetaldehyde (BAAA), a non-toxic fluorogenic substrate, into a fluorescent product by ALDH enzymes. Efflux of the fluorescent product was blocked by the efflux inhibitor provided in the Aldefluor buffer (Stemcell, cat. #01701), ensuring that fluorescence intensity reflected intracellular ALDH activity.

Briefly, 4 × 10 A549 cells were treated with 1 µM AB8939 for 3 hours at 37°C in a humidified 5% CO₂ incubator. Cells were then incubated with BAAA for 40 minutes at 37 °C in the presence of the efflux inhibitor, washed, and counterstained with DAPI to assess viability. Fluorescence was immediately analyzed by flow cytometry on an LSR II cytometer (BD Biosciences), and data were processed using DIVA software.

#### Mouse Experiments: MOLM-14 Xenograft, TG-LAM-75, and TG-LAM-36 PDX Models

All animal experiments were conducted in accordance with French regulations for animal care, approved by institutional ethics committees, and performed under sterile conditions. NOD.Cg-Prkdc^scid/J (NSG) mice, 6–10 weeks old, were maintained with sterilized food and water ad libitum on a 12 h light/dark cycle.

#### Cell Line–Derived Xenografts (MOLM-14)

Mice were injected via the tail vein with 2 × 10LJ viable luciferase-expressing MOLM-14 cells in 100 µL sterile PBS. Treatments began one day post-injection (D+1). Control animals received vehicle (20% hydroxypropyl-β-cyclodextrin in water). AB8939 was administered subcutaneously (SC) once daily at 1, 3, or 6 mg/kg, either 3 days ON/4 days OFF or 6 days ON/1 day OFF, starting at D+1. Vehicle was given SC 6 days ON/1 day OFF.

Mice treated with 1 mg/kg AB8939 or vehicle were sacrificed between days 24–30 due to leukemia progression. The 3 mg/kg cohort was terminated on day 33 for ethical reasons, with subsequent studies focused on 6 mg/kg dosing. Treatment at 6 mg/kg (3 days ON/4 days OFF) continued through six cycles (D37), after which disease progression required sacrifice. The 6 mg/kg (6 days ON/1 day OFF) cohort was treated until D53.

Mice were monitored daily for health and weight. Peripheral blood was collected by retro-orbital bleeding, and AML blasts were quantified by flow cytometry using PE anti-human CD33 (BD Pharmingen) and Pacific Blue anti-human CD45 (Ozyme). Bioluminescence imaging (BLI) was performed weekly with a PhotonIMAGER Optima (Biospace Lab) following anesthesia and intraperitoneal injection of endotoxin-free luciferin (30 mg/kg, Promega).

#### Patient-Derived Xenografts (TG-LAM-75, TG-LAM-36)

NSG mice were injected via the tail vein with 1 × 10LJ AML cells in 100 µL PBS. Upon detection of human CD33⁺ AML cells in peripheral blood by flow cytometry, mice were treated with:

- Vehicle
- Vidaza (2 mg/kg, 13 consecutive days, IP)
- Venetoclax (100 mg/kg, 7 consecutive days, oral)
- Cytarabine (Ara-C, 10 mg/kg, twice daily for 4 days, IP)
- AB8939 (6 mg/kg, 5 days ON/2 days OFF, SC)
- Vidaza + AB8939
- Venetoclax + AB8939

Disease progression was monitored by hCD33⁺ cells in peripheral blood, and tumor burden was quantified post-mortem in bone marrow and spleen. Investigators were not blinded to treatment allocation but were blinded during outcome assessment. Limiting dilution assays were performed in the TG-LAM-36 model to evaluate residual leukemia-initiating cells (LICs) after treatment (CTRL, Ara-C, AB8939). Mice showing BLI signal, hCD45⁺ cells in blood, or clinical signs of disease were considered leukemic.

#### AMKL Xenograft Model

The collection and use of human patient samples in this study was approved by the Internal Review Board of Gustave Roussy, with the informed consent of the patients in accordance with good clinical practice rules and national ethics recommendations. Mice were maintained in pathogen-free conditions and all experiments were approved by the Gustave Roussy institute animal care and use committee (Comité National de Réflexion Ethique sur l’Expérimentation Animale no. 26, project 2012-017). Patient-derived xenograft protocols and AMKL luciferase models were previously described (9,10). AMKL26 PDX cells transduced with a luciferase/mCherry lentivirus (gift of A.L. Kung, Dana-Farber Cancer Institute) were injected intravenously (3 × 10LJ cells) into irradiated (1.5 Gy) 6–12-week-old female NSG mice (Jackson Laboratory, stock #005557). Mice were maintained at Gustave Roussy under approved protocols (CEEA 26 #2017122111548235_v2; APAFIS #43692-2023053116192480 v3).

Engraftment was confirmed by BLI two weeks post-injection. Mice were then treated for 3 weeks with vehicle or AB8939 (2 mg/kg i.v., 3 days ON/4 days OFF for 2 weeks; 5 mg/kg, 3 days ON/4 days OFF for 1 week). BLI was performed weekly (IVIS Spectrum), and bone marrow was analyzed for human leukemic blasts 7 weeks post-injection.

#### Flow Cytometry

At week 7, bone marrow cells were flushed from femurs and tibias, stained in PBS/2% FBS for 30 minutes at 4 °C in the dark, washed, and analyzed by flow cytometry. mCherry⁺ human blasts were quantified by gating on live cells (Sytox Blue exclusion, Invitrogen). Data were acquired on FACSCanto II or FACSCanto X (BD) and analyzed with FlowJo software.

## Statistical Analysis

BLI data were expressed as mean ± SEM and analyzed by two-way ANOVA with Bonferroni post-tests. Day 22 BLI comparisons were performed using one-way ANOVA (Kruskal–Wallis) with Dunn’s multiple comparison test. Median survival was compared using the log-rank (Mantel–Cox) test. P-values <0.05 were considered significant. For PDX studies, one-way ANOVA analyses were performed using GraphPad Prism v10.6.1.

## Supporting information

Limiting Dilution Analysis

## Acknowledgments

The authors would like to thank the Institut Paoli-Calmettes Biological Resource Center and the Clinical Research and Innovation Department for their valuable contribution. We thank Pascal Verdier-Pinard (CRCM, Marseille, France) for having initiated in vitro cell-free microtubules polymerisation assays and *in cellulo* microtubules sedimentation analysis following AB8939 treatment. We are grateful to the staff of the CRCM animal facility for taking care of the mouse strain colonies. Diffraction data were collected at SOLEIL synchrotron (PROXIMA-1 and PROXIMA-2A beamlines, Saint-Aubin, France). We are most grateful to the machine and beam line groups for making these studies possible.

