## Supplementary figures and images for "Identification of AB8939, a novel synthetic microtubule destabilizer and ALDH inhibitor that overcomes multidrug resistance in tumor cells as a drug candidate for the treatment of refractory acute myeloid leukemia"

### Figure S1

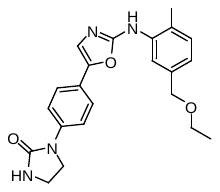

***Figure S1: Chemical structure of AB8939***

### Figure S2

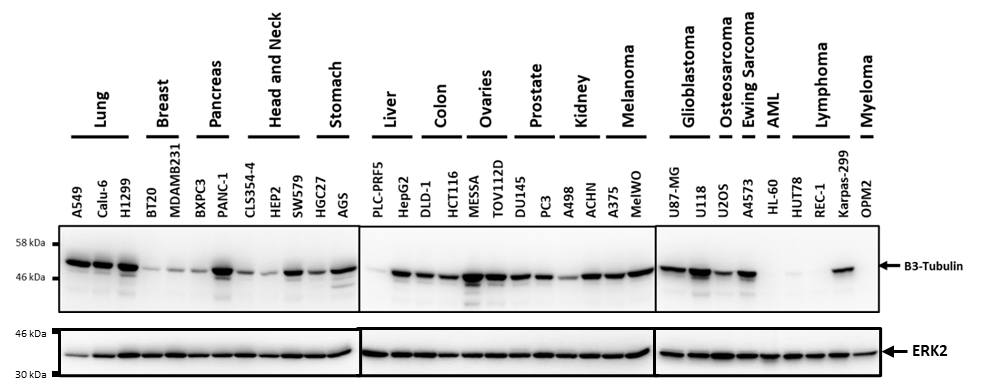

***Figure S2: β3-tubulin expression in tumor cell lines***

### Figure S4

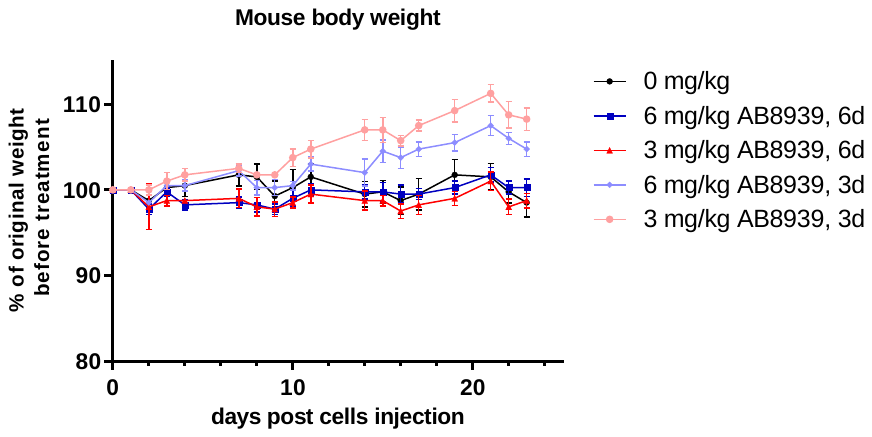

***Figure S4: Body weights monitoring over time for the 3 and 6mg/kg cohorts***
