## Supplementary material for "Identification of AB8939, a novel synthetic microtubule destabilizer and ALDH inhibitor that overcomes multidrug resistance in tumor cells as a drug candidate for the treatment of refractory acute myeloid leukemia": Figure S3

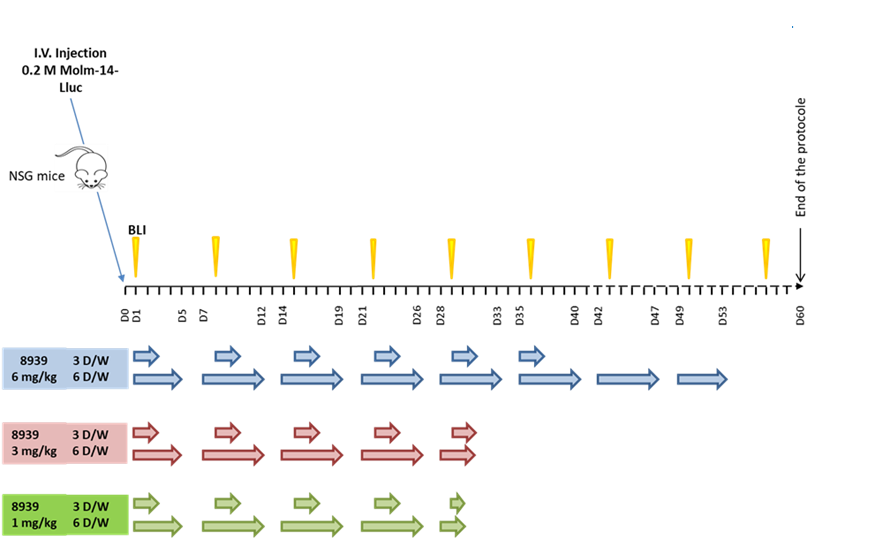

***Figure S3: Treatment protocol of NSG mice engrafted with MOLM14Lluc cell line, administration SC of AB8939 at 1, 3 or 6mg/kg, 3 or 6 days a week (3 or 6 D/W)***
