## Supplementary material for "Identification of AB8939, a novel synthetic microtubule destabilizer and ALDH inhibitor that overcomes multidrug resistance in tumor cells as a drug candidate for the treatment of refractory acute myeloid leukemia": Figure S5

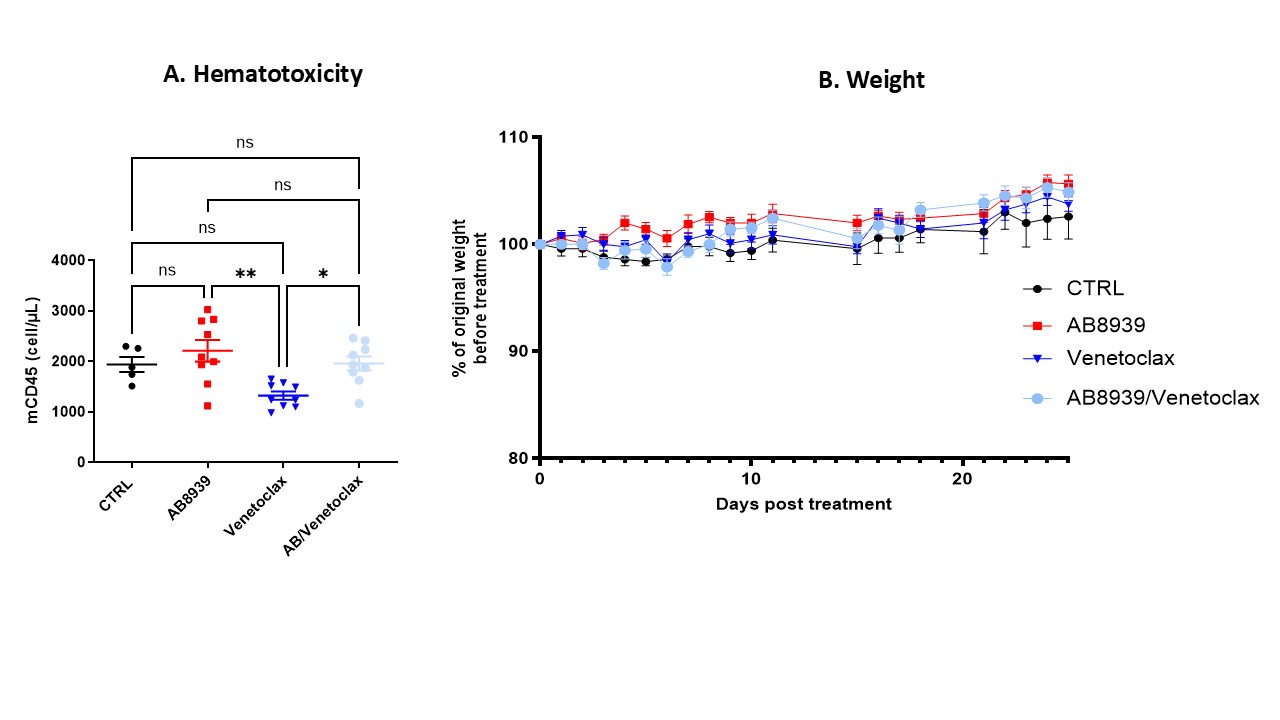
***Figure S5. Safety of AB8939 with venetoclax. Panel A: Murine CD45^+^ leukocyte quantification in peripheral blood (Day 31 post-engraftment, 14 days post-treatment initiation). Venetoclax, AB8939 and combination treatment preserved murine leukocyte counts at levels comparable to vehicle control, indicating absence of hematotoxicity. Data are presented as absolute cell counts (mCD45^+^ cells/µL blood) with individual values and mean ± SEM (vehicle n=5; AB8939 n=9; venetoclax n=9; combination n=9). Statistical significance determined by one-way ANOVA (***p < 0.001 azacitidine vs. Vehicle and AB8939). Panel B: Body weight monitoring over time for vehicle control, AB8939, venetoclax, and AB8939 plus venetoclax groups. Body weights are expressed as percentage of initial body weight. All treatment groups maintained stable weights throughout the study period, with no significant weight loss observed. This demonstrates that venetoclax, both alone and in combination with AB8939, is well tolerated without additional toxicity. All measurements were performed by flow cytometry. Data are presented as individual values with mean ± SEM. Statistical analyses performed using one-way ANOVA and Mann-Whitney t-test. While not statistically significant, the trends observed support further evaluation of the AB8939/venetoclax combination, and potentially a triplet regimen combining AB8939, azacitidine, and venetoclax.***
