## Supplementary material for "Identification of AB8939, a novel synthetic microtubule destabilizer and ALDH inhibitor that overcomes multidrug resistance in tumor cells as a drug candidate for the treatment of refractory acute myeloid leukemia": Limiting Dilution Analysis

**Limiting Dilution Analysis - Statistical Results**

LIC frequency comparison in experimental groups was calculated according to “the Extreme Limiting Dilution software from the Walter and Eliza Hall Bioinformatics Institute of Medical Research”; http://bioinf.wehi.edu.au/software/elda/ ).

**Table S4. Stem Cell Frequencies**

| Group | Frequency (1 in X cells) | 95% CI Lower | 95% CI Upper |
| --- | --- | --- | --- |
| CONTROL | 2,739 | 891 | 8,420 |
| ARAC | 4,193 | 1,396 | 12,592 |
| AB8939 | 32,181 | 4,622 | 224,072 |
| COMBO | 32,181 | 4,622 | 224,072 |

**Table S5. Statistical Analysis**

| **Overall Test for Group Differences** | | |
| --- | --- | --- |
| Chi-square | DF | P-value |
| 10.4 | 3 | 0.0156* |
| **Pairwise Comparisons** | | |
| Group 1 | Group 2 | Chi-square |
| AB8939 | ARAC | 3.96 |
| AB8939 | COMBO | 0.00 |
| AB8939 | CONTROL | 6.3 |
| ARAC | COMBO | 3.96 |
| ARAC | CONTROL | 0.271 |
| COMBO | CONTROL | 6.3 |

| Group 1 | Group 2 | Chi-square | P-value |
| --- | --- | --- | --- |
| AB8939 | ARAC | 3.96 | 0.0466* |
| AB8939 | COMBO | 0.00 | 1.000 |
| AB8939 | CONTROL | 6.3 | 0.0121* |
| ARAC | COMBO | 3.96 | 0.0466* |
| ARAC | CONTROL | 0.271 | 0.603 |
| COMBO | CONTROL | 6.3 | 0.0121* |

**P < 0.05 (statistically significant). Green highlighting indicates significant differences. DF = degrees of freedom.* Key Findings: 1. AB8939 and COMBO treatments significantly reduced stem cell frequency (~12-fold) compared to CONTROL (p = 0.0121); 2. ARAC alone showed no significant effect on stem cell frequency compared to CONTROL (p = 0.603); 3. AB8939 and COMBO showed identical frequencies (p = 1.0), indicating no additional benefit from combination therapy; 4. AB8939 appears to be the primary active agent responsible for stem cell depletion.
