## Supplementary material for "Identification of AB8939, a novel synthetic microtubule destabilizer and ALDH inhibitor that overcomes multidrug resistance in tumor cells as a drug candidate for the treatment of refractory acute myeloid leukemia": Table S1

***Table S1: Anti-proliferative activity of AB8939 and 8 MTAs on 32 tumor cell lines, in a 6-day proliferation/survival assay (IC_50_ nM).***

| Cell lines  IC_50_ (nM) | β3 Tubulin  (WB analysis) | AB8939 | Colchicine | CA-4 | Sabizabulin  (Veru-111) | Unesbulin  (PTC-596) | Lexibulin  (CYT997) | Vincristine | Eribulin | Taxol |
| --- | --- | --- | --- | --- | --- | --- | --- | --- | --- | --- |
| HL-60 | - | 14 | 10 | 2 | 7 | 48 | 8 | 1 | 2 | 0,3 |
| MOLM-14 |  | 14 |  | 12 |  |  | 45 |  |  |  |
| NOMO-1 |  | 46 |  | 94 |  |  | 280 |  |  |  |
| THP-1 |  | 49 |  |  |  |  | 9 |  |  |  |
| HUT-78 | -/+ | 4 | 7 | 1 | 3 | 20 | 6 | 2 | 1 | 2 |
| CCRF-CEM |  | 1 | 3 | 1 | 3 | 27 | 8 | 0,4 | 0,5 | 0,3 |
| Karpas-299 | ++ | 1 | 4 | 1 | 3 | 34 | 4 | 0,4 | 1 | 1 |
| REC-1 | -/+ | 4 | 19 | 1 | 2 | 17 | 10 | 56 | 125 | 346 |
| NALM-6 |  | 1 | 6 | 2 | 8 | 30 | 9 | 1 | 2 | 0,3 |
| A-549 | +++ | 1 | 25 | 4 | 6 | 13 | 21 | 10 | 3 | 0,3 |
| H-1299 | +++ | 0,3 | 3 | 2 | 1 | 5 | 7 | 2 | 75 | 0,3 |
| HEP-2 | + | 1 | 7 | 1 | 3 | 10 | 16 | 2 | 39 | 3 |
| HGC-27 | ++ | 1 | 4 | 2 | 2 | 9 | 8 | 1 | 0,3 | 0,3 |
| HRT-18 |  | 1 | 25 | 4 | 5 | 11 | 31 | 13 | 99 | 139 |
| PANC-1 | ++++ | 1 | 8 | 3 | 13 | 21 | 13 | 7 | 0,3 | 0,3 |
| PLC-PRF5 | +++ | 2 | 3 | 3 | 7 | 25 | 25 | 2 | 0,3 | 0,3 |
| U-118 | +++ | 1 | 4 | 3 | 4 | 33 | 13 | 0,3 | 0,3 | 0,3 |
| U87-MG | ++ | 0,3 | 4 | 2 | 4 | 11 | 20 | 1 | 0,3 | 0,3 |
| Tov21G |  | 8 | 12 | 0,4 | 61 |  | 13 | 1 |  |  |
| 786-O |  | 87 | 86 | 76 | 4 |  | 165 | 0,3 |  |  |
| ACHN | +++ | 4 | 15 | 2 | 4 |  | 36 | 166 |  |  |
| PC-3 | + | 8 | 14 | 63 | 1 |  | 142 | 23 |  |  |
| LnCaP |  | 5 | 3 | 14 | 4 |  | 52 | 66 |  |  |
| DU-145 | ++ | 2 | 32 | 3 | 8 | 17 | 32 | 4 | 2 | 0,3 |
| MCF-7 |  | 122 | 5 | 2 | 8 |  | 12 | 37 |  |  |
| MDA-MB-468 |  | 14 | 3 | 288 | 3 |  | 685 | 209 |  |  |
| SW-872 |  | 1 |  |  | 27 | 18 |  | 31 | 83 | 9 |
| SK-N-MC |  | 1 |  |  | 4 | 14 |  | 0,3 | 0,3 | 0,3 |
| A4573 | +++ | 3 | 3 |  | 3 | 21 |  | 1 | 0,3 | 0,3 |
| U2OS | ++ | 1 | 5 |  | 2 | 12 |  | 4 | 32 | 0,3 |
| WEHI-164 |  |  |  |  | 7 | 31 |  |  |  | 40 |
| MESSA | +++ | 7 | 10 | 2 |  |  | 24 | 106 | 121 | 10 |
| Average IC50 (nM) |  |  |  |  |  |  |  |  |  |  |
| Hemato |  | 15 | 8 | 14 | 5 | 29 | 42 | 10 | 22 | 58 |
| All Cells |  | 13 | 12 | 23 | 7 | 20 | 63 | 27 | 28 | 25 |
