## Supplementary material for "Identification of AB8939, a novel synthetic microtubule destabilizer and ALDH inhibitor that overcomes multidrug resistance in tumor cells as a drug candidate for the treatment of refractory acute myeloid leukemia": Table S2

***Table S2: Data collection and refinement statistics***

| Data collection^a^ |  |
| --- | --- |
| Space group | P2_1_2_1_2_1_ |
| Cell dimensions |  |
| a, b, c (Å) | 65.69, 128.76, 253.20 |
| *α, β, γ* (°) | 90.0, 90.0, 90.0 |
| Resolution (Å) | 126 – 2.53 (2.82 – 2.53) |
| Anisotropy resolution limits ^b^  R_meas_ | 3.31, 2.52, 2.82  0.145 (1.915) |
| I / σI | 12.7 (1.5) |
| CC_1/2_ | 0.996 (0.354) |
| Completeness (spherical)  Completeness (ellipsoidal) | 0.665 (0.127)  0.940 (0.688) |
| Multiplicity | 13.1 (13.1) |
| Refinement |  |
| Resolution (Å) | 126 – 2.53 |
| No. reflections | 47709 |
| Rwork / Rfree | 0.197 / 0.235 |
| No. atoms |  |
| Protein | 14295 |
| Ligands | 195 |
| Solvent | 55 |
| B factors |  |
| Protein | 94.4 |
| Ligands | 76.3 |
| Solvent | 63.7 |
| Coordinate error (Å) | 0.35 |
| R.m.s.d. |  |
| Bond lengths (Å) | 0.008 |
| Bond angles (°) | 0.90 |
| Ramachandran (%) |  |
| Favored region | 97.69 |
| Allowed region | 2.09 |
| Outliers | 0.22 |

^a^ Data were collected on a single crystal. Values in parentheses are for the highest-resolution shell. ^b^ Determined by STARANISO using a local mean I / σ(I)=1.2
