## Supplementary material for "Identification of AB8939, a novel synthetic microtubule destabilizer and ALDH inhibitor that overcomes multidrug resistance in tumor cells as a drug candidate for the treatment of refractory acute myeloid leukemia": Table S3

***Table S3. Comparative antiproliferative activity of AB8939, AraC, Azacytidine and Vincristine on AML patient cells coming from refractory-relapse CEGAL cohort.***

| AML Sample | | Genetics | | IC_50_ (µM) | | | |
| --- | --- | --- | --- | --- | --- | --- | --- |
| Inclusion_ID | **Class** | **karyotype** | **Mutations NGS** | **AB8939** | **AraC** | **AzaC** | **Vinc** |
| TG-LAM-76-P3 | M4 | 46,XY,inv(3)(q21q26)[20] | no mutation | 0,013 | 7,9 | 9,7 | nd |
| TG-LAM-136-P0 | M1 | 52,XY,+8,+10,+15,+3mar[20] | DNMT3A, IDH2, PHF6, NF1 | 0,04 | 13,1 | >50 | 1,35 |
| TG-LAM-75-P3 | M1 | 45,XX,**inv(3)(p23q26)**,**-7**[16] / 45,XX,sl,t(1;5)(q25;p15)[1] /45,sl,t(1;9)(q25;q34)[1] / 45,sl,t(1;11)(p25;p12)[1] / 45,sl,t(1;21)(q25;q22)[1] | TET2, **ASXL1**, RAS, RUNx1, IKZF1, BARD 1 | 0,05 | 4,1 | 13 | nd |
| TG-LAM-140-P0 | M5 | nd | **TP53**, NOTCH2, PTPN11, SF3B1 | 0,06 | >20 | 16,70 | 1,13 |
| TG-LAM-134-P0 | mixed | 46,XX,add(1)(p36),inv(9)(p12q12)?c,t(10;11)(p12-13;q14-21)[4] /46,sl,add(3)(q13),-15,-17,+2mar[16] | IKZF1, PTPN11, EZH2 | 0,08 | >20 | 39,10 | 0,17 |
| TG-LAM-141-P0 | LMMC2 | 46,X4,del(7)(q11q36)[25] | JAK2, **ASXL1**, U2AF1, **TP53**, Rad21, NRAS, ETV6 | 0,12 | 2,3 | 20,4 | 1,38 |
| TG-LAM-143-P0 | M4 | 47,XY,+8[5]/46,XY[16] | FLT3, SRSF2, TET2, NF1, **ASXL1**, CREBBP, RUNX1, MYD88 | 0,36 | 3,2 | 29,2 | 2 |
| TG-LAM-144-P0 | nd | 45,XX,del(7)(q21q31),der(7;17)(p10;q10),add(9)(q11),-21,+mar[20] | FLT3, APC, **TP53** | 0,38 | 11,5 | 41,7 | 1,605 |
| TG-LAM-139-P0 | M6 | 46,XX[20] | WT1, NPM1 | 0,94 | >20 | 38,50 | >2 |
| TG-LAM-145-P0 | MRC | 45,XY,add(4)(q35),del(5)(q15q34-35),+8,add(12)(q12-13),-16,-17,-19,+22[7] /45,sl,del(1)(q23q42)[11] / 45,sl,add(3)(q12),-3,-add(12),+12,-13,del(13)(q13q21), -15,add(20)(q12),+21,+19,+mar[2] /44,sl,-18,add(22)(p11)[2] | **TP53**, DNMT3A | 1,8 | >20 | 49,5 | >2 |
| TG-LAM-135-P0 | LAM M5 | 46,XY[20] | DNMT3A, TET2, NPM1 | >5 | >20 | >50 | >2 |
| TG-LAM-77-P0 | LAM M4 | 46,XX,del(9)(q13q33),?del(16)(q13q23)[19] / 46,XX[1] | JAK2, RUNX1 | >5 | >20 | 19,20 | >2 |
| TG-LAM-138-P0 | LAM M6 MRC | 46,XY[20] | SFSRF2, IDH2 | >5 | >20 | >50 | >2 |
| TG-LAM-142-P0 | LAM M2 | 47,XX,+8[21] / 46,XX[1] | GATA2, SRSF2, TET2, SUZ12 | >5 | >20 | >50 | >2 |
